# Recruitment of intrasegmental and propriospinal interneuron networks of forelimb motor circuitry in chronic cervical contusion spinal cord injury

**DOI:** 10.64898/2026.11.02.763629

**Authors:** Kallon Crowther, Angela Harbeck, Mackenzie Middlesworth, Morgan Bosch, Megan A. Lyttle, Claire MacGibbon, David A. Jaffe, Pauline Michel-Flutot, Evan Fitchett, Shuxin Li, Angelo C. Lepore

## Abstract

Cervical spinal cord injury (SCI) disrupts descending supraspinal input and spinal networks controlling upper extremity motor function, making recovery of arm and hand function a major priority for affected individuals. The vast majority of regenerative strategies have been studied during the acute and sub-acute phases following SCI, while therapeutic testing remains limited for chronic injury, when persistent barriers significantly limit circuit remodeling. We asked whether pharmacologic enhancement of neuronal-intrinsic axon growth capacity through PTEN inhibition could promote forelimb motor recovery and circuit plasticity in a rodent model of chronic cervical SCI. Following C5 hemi-contusion, female rats underwent behavioral testing of forelimb motor function and received PTEN antagonist peptide (PAP) beginning at a chronic time point, and the trans-synaptic tracer, pseudorabies virus (PRV), was injected into ipsilesional triceps to trace forelimb-associated motor circuitry. PAP selectively improved forelimb usage during cylinder paw-preference testing, but not strength, sensorimotor, or gait-related forelimb measures. PAP increased recruitment of PRV-traced local-projecting and propriospinal interneuron networks. In addition, PAP enhanced synaptic input to these recruited propriospinal interneurons, raising the possibility that spared descending pathways may contribute to recovery through spinal relay circuits. Lastly, we mapped the supraspinal connectome providing descending input from brainstem to the lower motor neurons innervating forelimb muscle, and we determined how this connectome was impacted both by SCI and PAP. Together, these findings show that, even in chronic SCI, pharmacological PTEN inhibition can promote forelimb functional recovery and significant circuit remodeling, with spinal interneuron networks emerging as a key site of PAP-induced plasticity.

**SIGNIFICANCE STATEMENT:** This study demonstrates that enhancing axon growth capacity in chronic cervical SCI using a non-invasive pharmacological approach can promote forelimb motor recovery and substantial remodeling of spared spinal circuitry. This is especially important because a vast majority of individuals with SCI are in the chronic phase, when treatment options are limited, and recovery of arm and hand function is highly desired. We also provide the first characterization of both spinal interneuron networks and supraspinal brainstem populations connected with triceps motor circuitry. Importantly, our findings identify local-projecting and propriospinal interneurons as key sites of treatment-induced plasticity. Together, this work expands our understanding of forelimb motor circuitry and highlights spinal relay networks as an important therapeutic target in chronic cervical SCI.

## INTRODUCTION

Spinal cord injury (SCI) can result in permanent deficits in autonomic, sensory, and motor functions (McDonald and Sadowsky, 2002; Ahuja et al., 2017). SCIs most often occur at the cervical level, disrupting critical motor systems that control the upper extremities and leading to long-lasting impairment (Venkatesh et al., 2019; Barbiellini Amidei et al., 2022). Restoration of arm and hand function is among the top priorities for patients with quadriplegia (Anderson, 2004). To date, several therapies have shown benefit for upper limb function when delivered at acute or subacute time points after injury (Walker et al., 2012; James et al., 2015). However, the vast majority of individuals living with SCI are in the chronic phase (Tashiro et al., 2017). In this late stage, injured spinal cord tissue develops a robustly inhibitory environment with extracellular barriers and reduced neuronal-intrinsic growth capacity, making therapeutic intervention particularly challenging (Tran et al., 2018). It is imperative to develop effective treatments that can overcome these inhibitory barriers and promote significant functional recovery in the context of chronic cervical SCI.

A single isolated pathway does not mediate forelimb motor control. It instead emerges from coordinated interactions across multiple descending supraspinal systems, spinal interneuron networks, afferent connections, and neuromodulatory input (Azim et al., 2014; Baker et al., 2015; Sheikh et al., 2018; Wang et al., 2018). Within the spinal cord, local-projecting and propriospinal interneurons serve as both intra- and intersegmental relays that shape motor output (Alstermark et al., 2007; Zholudeva et al., 2021). These interneuron populations are highly heterogeneous and are broadly distributed throughout the spinal gray matter (Zholudeva et al., 2021). They can project locally or across multiple spinal segments (Laliberte et al., 2019). Their widespread distribution and diverse connectivity allow them to relay descending and other inputs to lower motor neurons (LMNs) (Alstermark et al., 2007; Zholudeva et al., 2021). Descending motor pathways are often polysynaptic, transmitting supraspinal commands through these spinal networks, while descending neuromodulatory systems further refine spinal output (Baker et al., 2015; Wang et al., 2018; Giorgi et al., 2025). Together, highly distributed and interconnected systems controlling forelimb motor output provide a significant challenge in identifying which forms of circuit plasticity are the most relevant to achieving functional recovery after SCI.

One strategy to promote circuit plasticity after SCI is to enhance the growth capacity of injured or spared neurons. Adult central nervous system (CNS) neurons have limited intrinsic axon growth capacity, which is further restricted after CNS injury (Sun and He, 2010). A key regulator of neuron-intrinsic growth ability is phosphatase and tensin homolog (PTEN), a negative regulator of phosphatidylinositol 3-kinase (PI3K)/Akt/mammalian target of rapamycin (mTOR) signaling that is highly expressed in adult CNS neurons (Park et al., 2010; Hausott et al., 2022). Previous studies have shown that PTEN inhibition promotes axon growth and circuit reorganization in both spared and injured pathways, resulting in functional recovery in SCI models (Walker et al., 2012; Ohtake et al., 2014). Inducible genetic PTEN deletion in the chronically injured adult mouse CNS promotes corticospinal (CST) tract axon growth (Du et al., 2015). More recent work using AAV-mediated PTEN knockout three months after SCI demonstrated transient improvements in locomotor function that varied with injury severity (Stewart et al., 2023). We previously developed a PTEN antagonist peptide (PAP), a blood-brain barrier-penetrant peptide that enables systemically administered pharmacologic inhibition of PTEN within the CNS (Ohtake et al., 2014). In multiple models of chronic cervical SCI, we have demonstrated that PAP can promote bulbospinal axon growth and partial recovery of diaphragmatic respiratory motor function (Cheng et al., 2021; Michel-Flutot et al., 2024). These findings suggest that chronically injured motor circuitry can respond to manipulation of neuron-intrinsic axon growth capacity. Whether pharmacologic PTEN inhibition in the context of chronic cervical SCI drives remodeling across the highly distributed networks controlling forelimb function remains unclear.

Cervical hemi-contusion models are widely used to study forelimb motor deficits and recovery after SCI because they partially preserve descending supraspinal and spinal interneuron pathways (Anderson et al., 2009; Mondello et al., 2015; Freria et al., 2023), providing substrates for post-injury circuit remodeling. However, little is known about how enhancing neuron-intrinsic axon growth capacity affects forelimb-associated circuitry in chronic cervical contusion SCI. In this study, we used a chronic cervical level-5 (C5) hemi-contusion SCI model and pharmacologic PTEN inhibition with PAP to examine forelimb motor recovery and remodeling across multiple components of the motor network. Our findings identify several forms of circuit plasticity that are associated with improved forelimb motor function in response to enhancement of axon growth capacity in chronic cervical SCI.

## MATERIALS AND METHODS

### Animals

We used female Sprague-Dawley rats (250-300 grams, Taconic Farm, Rockville, MD). Thomas Jefferson University Institutional Animal Care and Use Committee (IACUC) approved all experimental procedures and ensured they were conducted in compliance with the *National Institutes of Health (NIH) Guide for the Care and Use of Laboratory Animals* and the *ARRIVE (Animal Research: Reporting of In Vivo Experiments)* guidelines. In a temperature, humidity, and light-controlled environment, animals were housed three animals per cage. Food and water were provided *ad libitum* on a 12-hour light/dark cycle. In previous work, we have optimized and characterized the unilateral cervical contusion SCI model in adult female rats and the PAP delivery paradigm in chronic SCI (Nicaise et al., 2012; Nicaise et al., 2013; Michel-Flutot et al., 2024)

### Experimental design

All animals were randomized, and the experimental timeline is shown in Figure 1. Animals underwent pre-injury baseline behavioral testing, with repeated behavioral assessments throughout the experiment. Thirty animals underwent a left-sided C5 hemi-contusion. Beginning 4 weeks post-injury, animals received either DMSO-only (vehicle control) or PAP by subcutaneous injection every 12 hours for 6 weeks. PAP was administered at 1 mg/day/rat, as previously described (Urban et al., 2019). Behavioral testing continued through 18 weeks post-injury. At week 22, pseudorabies virus (PRV) was injected into the ipsilateral triceps muscle for trans-synaptic tracing. Animals were euthanized 4.5 days later, and tissue was collected for histological analysis. Four animals were excluded because of surgical inconsistencies or mortality during the C5 hemi-contusion procedure. Two additional animals were euthanized at different post-injury time points after reaching predefined humane endpoints due to contralateral hindlimb autophagy. One additional animal was excluded from all behavioral analyses because severe post-injury impairment prevented meaningful participation in the behavioral assays.

**Figure 1.**
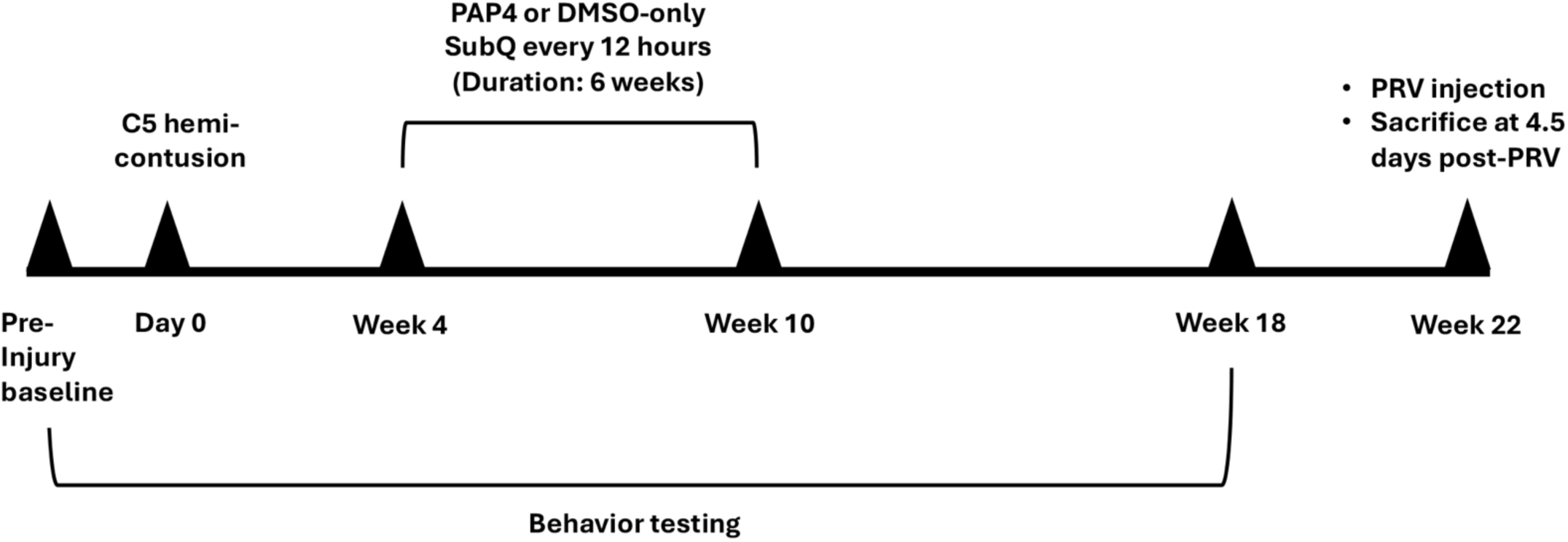
Experimental timeline. Rats underwent baseline behavioral testing followed by a unilateral C5 hemi-contusion on day 0. Beginning at 4 weeks post-injury, animals received either DMSO vehicle or PAP by subcutaneous injection every 12 h for 6 weeks. Behavioral assessments were performed through 18 weeks post-injury. At 22 weeks post-injury, rats received ipsilateral triceps injection of the trans-synaptic tracer PRV-152. Animals were euthanized 4.5 days after PRV inoculation for histological analysis.

### C5 unilateral cervical contusion

All injured animals received a left-sided hemi-contusion. Prior to anesthesia, animals received subcutaneous buprenorphine (Endo Pharmaceuticals Inc., Malvern, PA; 0.03 mg/kg), sulfamethoxazole-trimethoprim (Somerset Therapeutics, Somerset, NJ; 30 mg/kg), and carprofen (Zoetis Inc., Parsippany, NJ; 5 mg/kg). Anesthesia was induced with 5% isoflurane (Piramal Critical Care, Bethlehem, PA) in 100% O₂ in a closed chamber and maintained at 1.5–2% isoflurane in 100% O₂ throughout the procedure. Animals were placed on a heating pad to maintain temperature during the surgery. The skin and muscles were retracted to expose the spinal column. A laminectomy and durotomy were performed at the C5 level. Rats then underwent a left-sided C5 contusion using the Infinite Horizon Spinal Impactor (Precision Systems and Instrumentation, Lexington, KY) with a 1.5 mm tip and a target force of 200 kdyn, based on previously published cervical contusion parameters (Nicaise et al., 2012; Freria et al., 2023). The impactor was set to a velocity of 1 mm/s with a dwell time of 0 s. Following the contusion, muscles were sutured with sterile 4-0 Vicryl and the skin was closed with sterile wound clips (Braintree Scientific, Braintree, MA). Animals received lactated Ringer’s solution (4 mL, subcutaneously) and were monitored during recovery from anesthesia and throughout the postoperative period.

### Behavioral testing

Animals were habituated to handling for 14 consecutive days using an adapted protocol previously established in our laboratory (Deacon, 2006). Prior to baseline testing, animals received limited exposure to the behavioral apparatuses. Behavioral testing was conducted in the same room by the same handler throughout the study to reduce variability.

### Grip strength test

Forelimb grip strength was assessed using a Chatillon grip strength meter (AMETEK, Largo, FL). The left and right forelimbs were tested individually. For each trial, the animal grasped the meter, and the handler pulled consistently backward until the animal released its grip. Grip strength was recorded in grams of force. Three measurements were obtained for each forelimb at each testing time point, and the mean of the three trials was used for analysis.

### Grid sensorimotor test

Animals were placed on an elevated 42 x 42 cm wire grid positioned 42 cm above the tabletop, with 3.8 x 3.8 cm square openings. Animals were allowed to explore the grid freely for 2 min while behavior was recorded from below to permit visualization of individual paw placements. Left and right foot faults and left and right successful placements were scored post hoc from video recordings.

### Paw print gait test

Paw print analysis was adapted from previously published gait assessment methods (Dunham et al., 2010; Wertman et al., 2019). Animals traversed a narrow, walled paper runway approximately 152 cm in length and 11.5 cm in width toward a dark goal box. Prior to each trial, the left forepaw was coated with red non-toxic paint and the right forepaw with black non-toxic paint. Animals were allowed to traverse the runway freely. Trials were repeated if animals turned around, stopped forward progression, or otherwise failed to complete the runway. When repeat trials were required, animals were rested for at least 10 min between attempts. Stride length and toe spread were measured directly from paw prints using digital calipers. Stride length was defined as the distance between two consecutive placements of the same forepaw, and toe spread as the distance between the most medial and most lateral toes. Paw surface area was quantified from digitally scanned paw prints using ImageJ (RRID: SCR_003070). For each animal and testing time point, stride length and toe spread were averaged across three paw prints, whereas paw surface area was averaged across two paw prints.

### Cylinder paw-preference test

Animals were placed individually in a transparent cylindrical enclosure approximately 20 cm in diameter and 32 cm in height and allowed to explore freely for 5 min while behavior was recorded. Mirrors were positioned around the cylinder to allow visualization of forelimb usage when animals turned away from the camera. Right forelimb touches, left forelimb touches, and bilateral forelimb touches were scored post hoc from video recordings.

### Retrograde trans-synaptic tracing of forelimb motor circuitry

Four and a half days before euthanasia, animals were anesthetized with isoflurane (5% induction, 1.5–2% maintenance in 100% O₂). Before anesthesia, animals received subcutaneous EthiqaXR (Fidelis Animal Health, Inc., North Brunswick, NJ; 0.65 mg/kg), sulfamethoxazole-trimethoprim (Somerset Therapeutics, Somerset, NJ; 30 mg/kg), and carprofen (Zoetis Inc., Parsippany, NJ; 5 mg/kg). Animals were maintained on a heating pad throughout the procedure. The skin over the left upper forelimb was gently retracted to expose the triceps muscle, and loose fascia was removed. PRV-152 (PRV-Bartha; Princeton Neuroscience Institute Viral Core Facility; 4.13E9 pfu/mL) was injected at four sites along the length of the muscle for a total volume of 20 µL. The skin was closed with sterile wound clips, and animals received lactated Ringer’s solution (4 mL, subcutaneously). Animals were monitored during recovery from anesthesia and throughout the postoperative period.

### Tissue processing

All animals were transcardially perfused with 1X phosphate-buffered saline (PBS), followed by 4% paraformaldehyde (PFA). The brain, brainstem, and spinal cord were carefully dissected and post-fixed in 4% PFA at 4°C for 24 hours. Tissue was then transferred to 30% sucrose in 1X PBS for cryoprotection for 48 hours. Frozen transverse spinal cord sections (30 µm) and brain sections (40 µm) were cut free-floating on a Microm HM550 cryostat (Thermo Fisher Scientific, Waltham, MA). Before sectioning, a small notch was made on the contralateral side of the spinal cord and on the ipsilateral side of the brain to preserve left-right orientation during free-floating processing. All sections were stored at −20°C in cryoprotectant solution containing 30% sucrose, 30% ethylene glycol, and 1% PVP-40 in 1X PBS.

### Cresyl violet staining and lesion analysis

Lesion area and volume were quantified as previously described in our laboratory (Lepore et al., 2008; Lepore et al., 2011a; Nicaise et al., 2012). Selected spinal cord sections were stained with 0.5% cresyl violet acetate to assess lesion morphology and quantify LMNs. Brightfield images were acquired using a Keyence BZ-X1000 microscope (KEYENCE Corporation of America, Itasca, IL). Lesion and spared tissue areas were quantified on every 12th section using MetaMorph Microscopy Automation and Image Analysis Software (Molecular Devices; RRID: SCR_002368) by manually tracing the total spinal cord and lesion boundaries. Lesion was defined as regions containing either tissue loss/cavitation or surrounding damaged tissue in which normal spinal cord architecture was disrupted, whereas spared tissue was defined as regions in which normal anatomical structure was preserved. Lesion volume was calculated using the Cavalieri estimator: V = [Σ (A₁ + A₂ + … + Aₙ) × D] − [Amax × Y], where D represents the distance between measurements and Y represents section thickness. Ventral horn LMN cell bodies were manually identified and counted in the same cresyl violet-stained sections. Motor neuron cell bodies with a clearly identifiable nucleolus and a soma of >200 μm^2^ were counted (Lepore et al., 2011b).

### Immunohistochemistry

Free-floating sections were washed three times for 5 min in 1X PBS before immunohistochemical processing. Unless otherwise specified, sections were blocked in 5% normal donkey serum (NDS) and 0.2% Triton X-100 in 1X PBS, followed by overnight incubation at 4°C in primary antibody solution containing 5% NDS and 0.1% Triton X-100 in 1X PBS. Primary antibodies included mouse anti-synaptophysin (1:250; Abcam, cat. no. ab8049; RRID: AB_2198854), chicken anti-GFP (1:2000; Aves Labs; RRID: AB_10000240), rabbit anti-5-HT (1:10,000; ImmunoStar; RRID: AB_572263), and guinea pig anti-IBA1 (1:500; Synaptic Systems, cat. no. 234308; RRID: AB_2924932). Anti-GFP immunolabeling was used to amplify endogenous PRV-152 GFP signal. For 5-HT immunostaining, sections were blocked for 2 h at room temperature in 5% NDS, 1% Triton X-100, and 0.05% Tween 20 in 1X PBS. For Iba1 immunostaining, normal goat serum was used in place of NDS. Following primary antibody incubation, sections were washed three times for 5 min in 1X PBS and incubated for 2 h at room temperature with the appropriate secondary antibodies diluted 1:2000 in 5% normal serum and 0.1% Triton X-100 in 1X PBS. Secondary antibodies included Alexa Fluor 488 donkey anti-chicken (Invitrogen, cat. no. A78948; RRID: AB_2921070), Alexa Fluor 594 donkey anti-rabbit (Invitrogen, cat. no. A21207; RRID: AB_141637), Alexa Fluor 647 donkey anti-mouse (Invitrogen, cat. no. A31571; RRID: AB_162542), and Alexa Fluor 594 goat anti-guinea pig (Invitrogen, cat. no. A11076; RRID: AB_2534120). Sections were washed three times for 5 min in 1X PBS, mounted onto slides, and cover-slipped using Fluoromount-G Mounting Medium with DAPI (Invitrogen, cat. no. 00-4959-52).

### Spinal cord PRV-labeled interneuron quantification

PRV-positive cells within the spinal cord were manually counted using a Zeiss Axio Imager M2 epifluorescence microscope (Carl Zeiss Microscopy, LLC, Thornwood, NY). Counts were performed separately within the dorsal horn, intermediate zone, and ventral horn. Three sections were analyzed at each of three distances from the lesion epicenter (1.44, 1.80, and 2.16 mm), both rostral and caudal to the injury, for a total of nine sections per direction per animal. Counts were summed across the three sections at each distance for analysis.

### Brainstem PRV-labeled neuron quantification

Brainstem sections were imaged using a Zeiss Axio Imager M2 epifluorescence microscope (Carl Zeiss Microscopy, LLC, Thornwood, NY). Images spanning the brainstem were manually stitched to generate composite images for analysis. Brainstem nuclei were manually delineated using anatomical landmarks and reference to the rat brain atlas (Paxinos and Watson, 2014). PRV-positive cells were manually counted within the caudal pontine nucleus (PnC), ventral pontine nucleus (PnV), gigantocellular reticular nucleus (Gi), gigantocellular reticular nucleus alpha (GiA), ventral gigantocellular reticular nucleus (GiV), lateral paragigantocellular nucleus (LPGi), lateral paragigantocellular nucleus alpha (LPGiA), raphe magnus (RMg), raphe obscurus (ROb), raphe pallidus (RPa), and the dorsal and ventral medullary reticular nuclei (MdD/MdV). MdD and MdV were combined for analysis. Counts were performed separately ipsilateral and contralateral to the lesion for bilateral nuclei, while RMg, ROb, and RPa were counted along the midline.

### Fluorescence intensity analysis

Iba1 immunofluorescence intensity was quantified in intact caudal and peri-lesional spinal cord sections using MetaMorph Microscopy Automation and Image Analysis Software (Molecular Devices; RRID: SCR_002368). Within each section, the dorsal horn, intermediate gray, and ventral horn were manually delineated as regions of interest using the tracing tool. Mean Iba1 fluorescence intensity within each region of interest was recorded and expressed in arbitrary units (A.U.). Image acquisition settings were held constant across all Iba1 images included in the analysis. For 5-HT analysis, fluorescence intensity was quantified within the ventral horn caudal to the lesion using sections selected from the previously defined caudal sampling region. The ventral horn was manually delineated as a region of interest using MetaMorph, and the mean 5-HT fluorescence intensity was recorded and expressed in arbitrary units. Image acquisition settings were held constant across all 5-HT images included in the analysis.

### Confocal imaging and synaptic analysis

Representative fluorescent images were acquired using a Leica SP8 confocal microscope (Leica Microsystems Inc., Buffalo Grove, IL) with LAS X imaging software (RRID: SCR_013673). For synaptophysin analysis, dorsal horn sections within the previously defined rostral sampling region used for PRV quantification were triple-labeled for PRV-GFP, synaptophysin, and 5-HT. Confocal z-stacks spanning 20 µm were acquired at 40× magnification using 10 optical steps. Approximately 10 PRV-positive cells were analyzed per animal. For each cell, synaptophysin-positive puncta associated with the PRV-GFP-positive cell body and 5-HT/synaptophysin double-positive puncta associated with the PRV-GFP cell body were quantified in a single section within the z-stack. Cell-level values were averaged within each animal to generate a single value per animal.

### Statistical analyses

Data are presented as mean ± SEM throughout. Details of sample sizes and results of all statistical analyses, including exact *p*-values and test statistics, are provided in the corresponding supplemental tables. Statistical analyses and graphing were performed using GraphPad Prism 10.6.1 (GraphPad Software, Boston, MA), with statistical significance defined as *p* < 0.05. Longitudinal behavioral outcomes and rostro-caudal histological analyses were analyzed using mixed-effects models. Post hoc comparisons were performed using Šídák multiple-comparisons tests where indicated. 5-HT fluorescence intensity was analyzed using one-way ANOVA followed by Tukey multiple-comparisons testing. Two-group comparisons, including brainstem and spinal cord PRV-positive cell counts, were analyzed using *t*-tests. Normality was assessed for each data set, and appropriate statistical tests were used based on this analysis.

### Data availability

All raw data from this study will be deposited in the Open Data Commons for Spinal Cord Injury (ODC-SCI) for open sharing, in accordance with FAIR data principles.

## RESULTS

### PAP treatment after chronic cervical contusion SCI promoted partial recovery of forelimb motor function

Adult rats subjected to a C5 hemi-contusion SCI received twice-daily systemic administration of PAP or DMSO-vehicle-only control starting at the chronic time point of four weeks post-injury and lasting for six weeks until week 10 (Figure 1). To assess effects on forelimb motor function, animals were tested using a battery of behavioral assays starting at pre-injury baseline and continuing until 18 weeks following contusion. Grip testing was used to determine forelimb muscle strength (Figure 2A). Grip strength of the contralateral forelimb was unaffected by the contusion (Figure 2C), while the ipsilateral forelimb showed a small degree of increased grip strength (Figure 2B), possibly due to SCI-induced spasticity. For both ipsilateral and contralateral forelimbs, there were no differences between PAP and DMSO-only groups at all time points (Figure 2B-C; statistical details for grip analysis provided in Supplemental Table 1). Grid walk testing was employed to examine sensorimotor function of each forelimb (Figure 2D). C5 contusion SCI resulted in a significant and persistent increase in faults during placement of the ipsilateral forepaw (Figure 2E), while the contralateral forepaw was unaffected by the SCI (Figure 2F). For both ipsilateral and contralateral forelimbs, there were no differences between PAP and DMSO-only at all assessed time points (Figure 2E-F; statistical details for grid analysis provided in Supplemental Table 1). Paw print analysis evaluated gait-related measures of forelimb usage during locomotion (Figure 2G-H). C5 contusion resulted in robust and persistent alterations in stride length (Figure 2I), but only limited and transient changes in other ipsilateral forepaw measures such as length of ipsilateral toe spread during paw placement (Figure 2J) and surface area (Figure 2K). Furthermore, no differences between groups were noted for any of the assessed parameters at any time point after injury (Figure 2I-K; statistical details for paw print analysis provided in Supplemental Table 1). Lastly, cylinder testing was performed to determine paw preference usage during rearing events as a measure of forelimb motor function (Figure 2L). Following C5 contusion, both DMSO-only and PAP animals showed a dramatic reduction in use of the ipsilateral forepaw during rearing (Figure 2M). Compared to DMSO-treated rats, PAP animals showed a significant and persistent recovery in use of the affected ipsilateral forelimb out to 18 weeks post-injury (treatment: *p* = 0.023; time *p* = 0.0002; treatment x time interaction: *p* = 0.017). Statistical details for cylinder analysis are provided in Supplemental Table 1. Together, these findings demonstrate that PAP treatment starting at a chronic time after cervical SCI partially improved forelimb motor function and that this effect was selective to only a subset of behavioral measurements.

**Figure 2.**
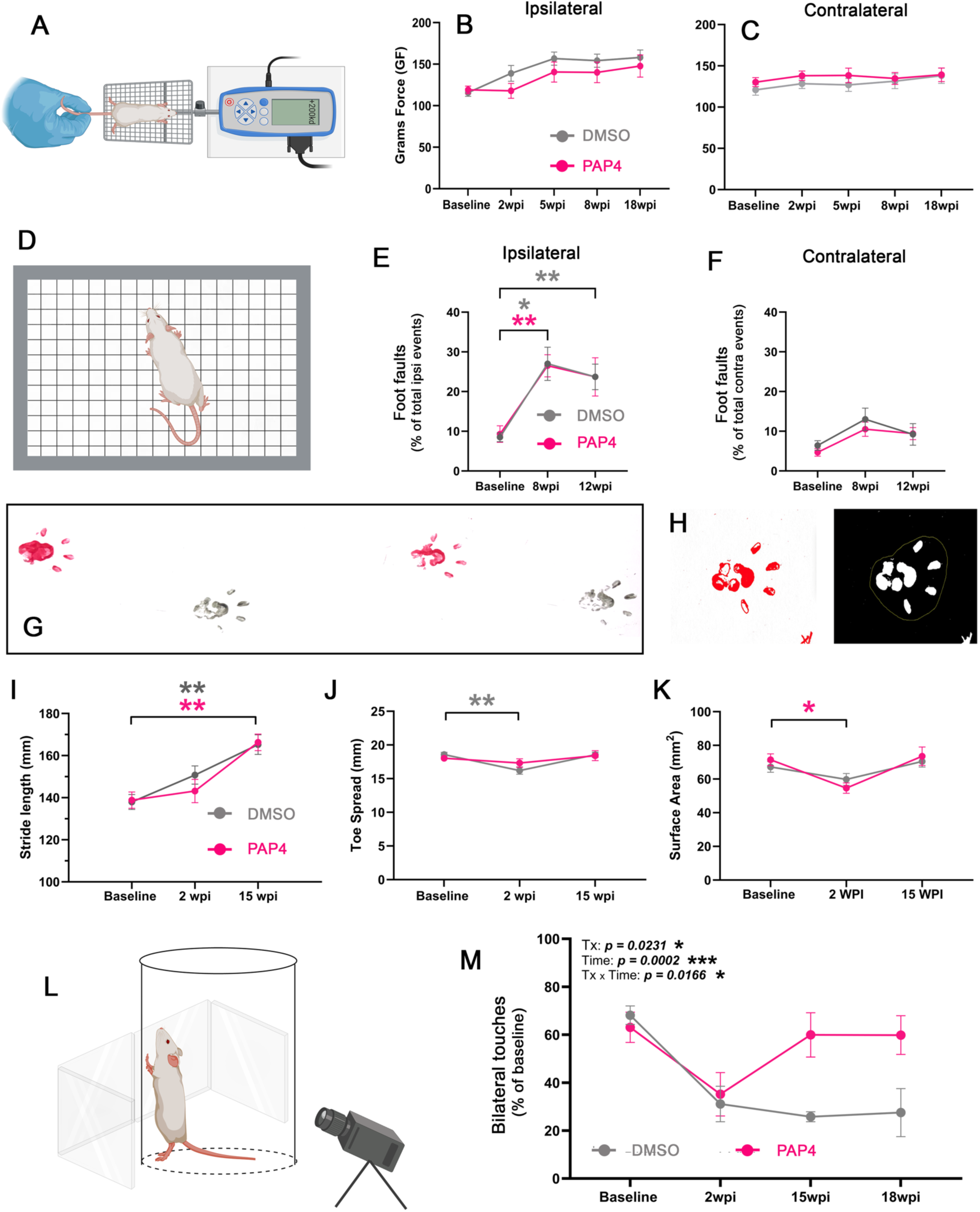
Behavioral assessment of forelimb motor function following chronic cervical contusion SCI and PAP treatment. Schematic of forelimb grip strength testing **(A).** Ipsilateral **(B)** and contralateral **(C)** grip strength across the experimental time course. Ipsilateral grip strength showed a significant time effect, with no significant treatment effect. Schematic of grid sensorimotor testing **(D).** Ipsilateral **(E)** and contralateral **(F)** foot faults across testing time points. Foot faults increased significantly over time on the ipsilateral side, with no significant treatment effects. Representative paw prints used for gait analysis **(G)** and paw surface-area quantification **(H).** Ipsilateral stride length **(I),** toe spread **(J),** and paw surface area **(K)** across the experimental time course. All three measures showed significant effects of time. Schematic of cylinder paw-preference testing **(L).** Bilateral forelimb contacts as a percentage of total forelimb contacts at baseline and 2, 15, and 18 weeks post-injury **(M).** Mixed-effects analysis showed significant effects of treatment, time, and treatment × time, with exact p-values shown on the graph. \**p* < 0.05; **p < 0.001. Details of statistical analyses are provided in Supplemental Table 1.

### PAP administration did not alter lesion size or neuronal degeneration following cervical SCI

Contusion-type SCI is accompanied by significant secondary expansion of the lesion and degeneration of neuronal populations following the initial traumatic insult. As behavioral testing demonstrated that PAP promoted partial recovery of forelimb motor function, we determined whether this improvement was associated with effects on preservation of cervical spinal cord tissue and protection of LMNs. Using quantitative histological analysis of cresyl violet-stained transverse sections of cervical spinal cord (Figure 3A-G), we found that PAP did not alter two-dimensional lesion area at the injury epicenter or in surrounding rostral and caudal locations compared to DMSO-only control (Figure 3H). In addition, there was no difference in overall three-dimensional lesion volume between DMSO-only and PAP conditions (Figure 3I; statistical details for lesion analysis provided in Supplemental Table 2). Furthermore, numbers of surviving ventral horn LMNs ipsilateral to the contusion site were quantified in cresyl violet-stained transverse sections (Figure 4A-C). There was no difference in LMN counts between DMSO-only and PAP across all rostral-caudal distances from the lesion epicenter (Figure 4D). Statistical details for LMN counts are provided in Supplemental Table 2. These lesion area/volume and LMN count analyses demonstrate that PAP did not alter tissue degeneration or neuronal loss following cervical contusion SCI. These findings also show consistency of the SCI across animals in this study.

**Figure 3.**
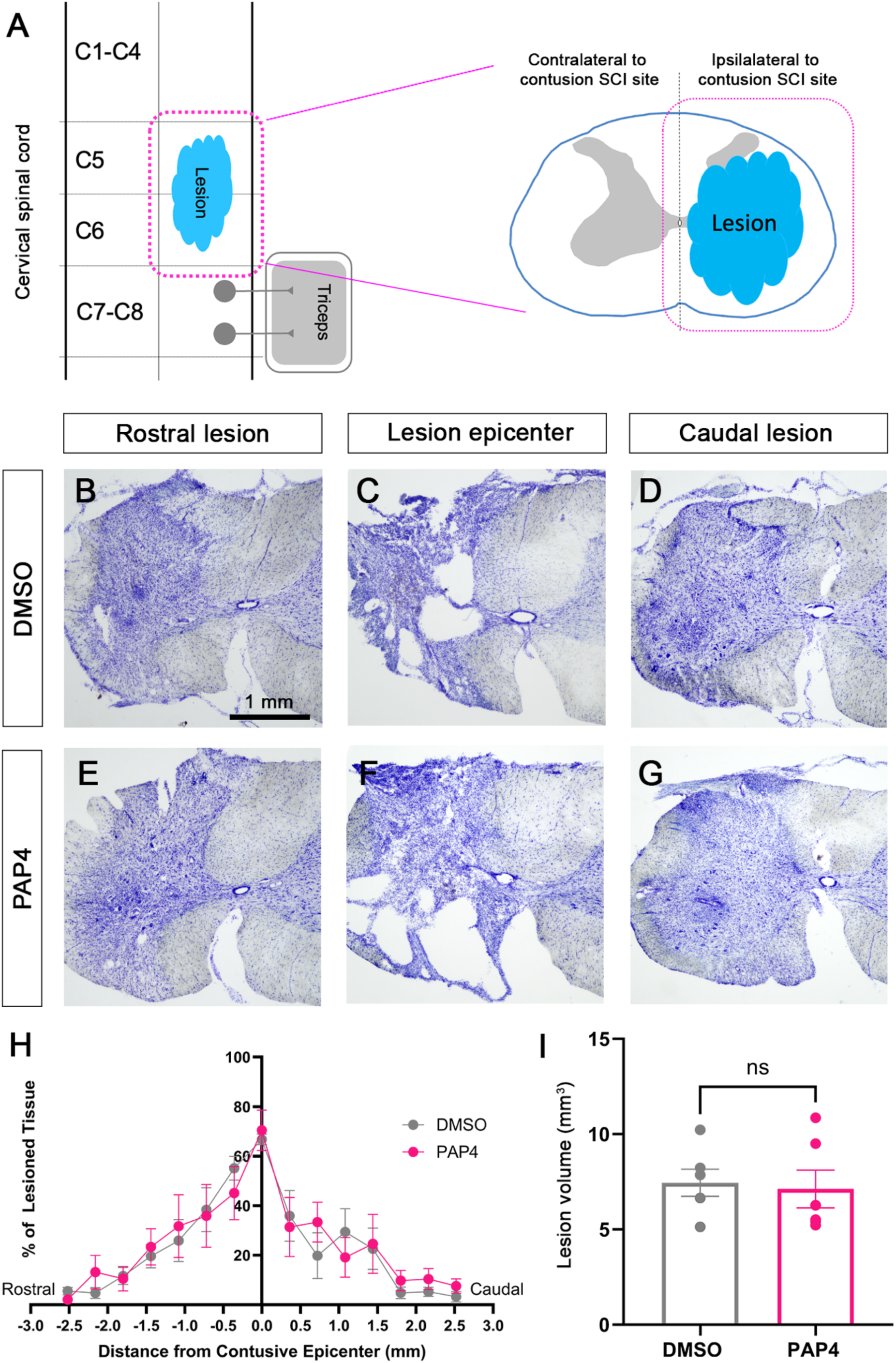
Lesion area and volume following chronic cervical contusion SCI. Schematic depicting unilateral cervical contusion SCI and a cross-section of the injured spinal cord **(A)**. Representative cresyl violet-stained spinal cord sections from DMSO-only animals in the rostral region of injury **(B)**, lesion epicenter **(C)**, and caudal lesion region **(D)**. Representative sections from PAP-treated animals are shown at the corresponding rostral **(E)**, epicenter **(F)**, and caudal **(G)** locations. Quantitative lesion analysis showed no significant differences in lesion area between treatment groups across the rostrocaudal extent of the injury **(H)**. Total lesion volume, calculated using the Cavalieri estimator, also did not significantly differ between groups (I). Details of statistical analyses are provided in Supplemental Table 2.

**Figure 4.**
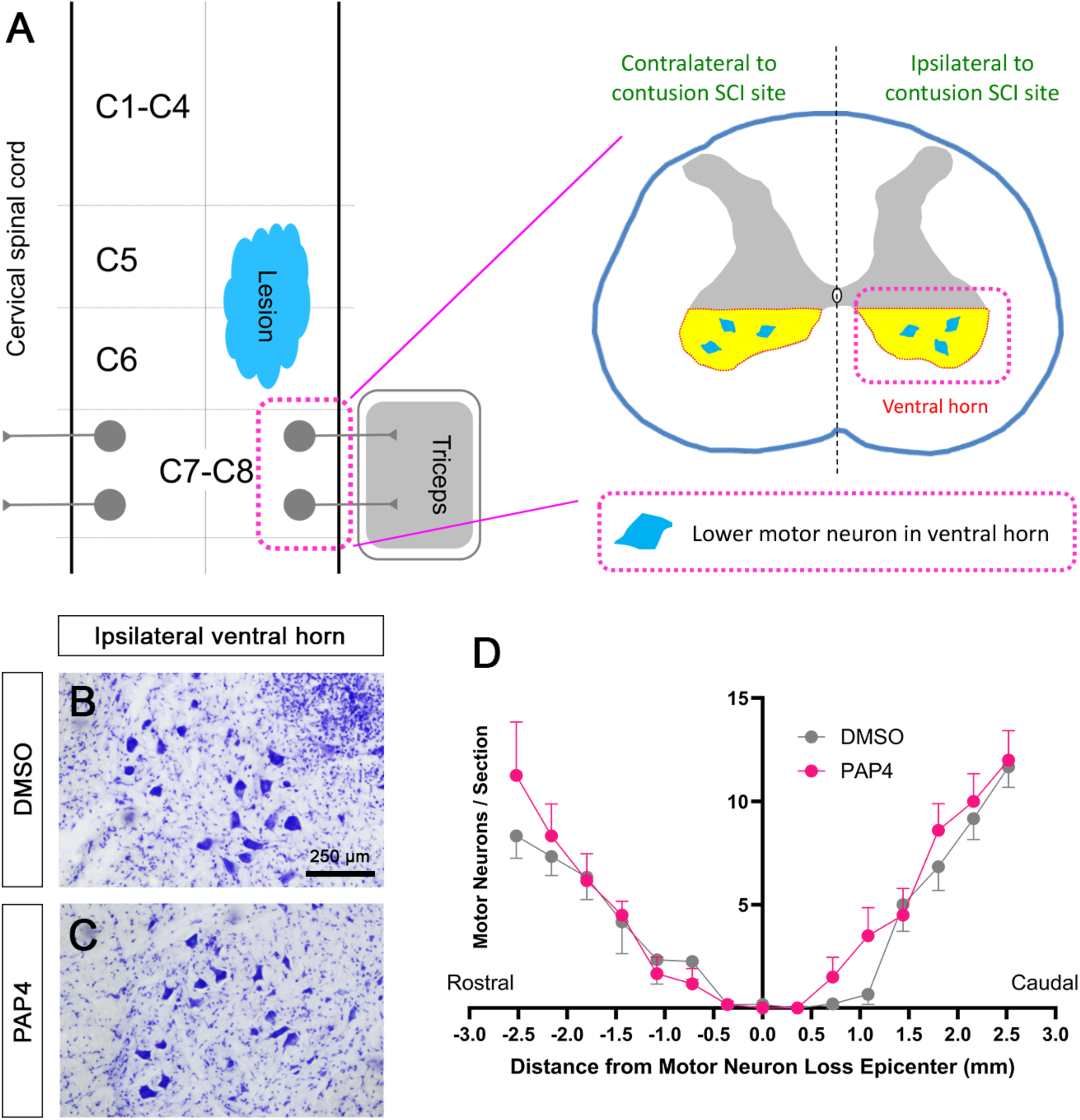
Ventral horn lower motor neuron quantification. Schematic illustrating unilateral cervical contusion SCI and the ipsilateral ventral horn region used for quantification **(A).** Representative cresyl violet-stained spinal cord sections from DMSO-only and PAP-treated animals depicting ventral horn lower motor neurons **(B–C).** Quantitative analysis of lower motor neuron counts showed no significant differences between treatment groups **(D).** Details of statistical analyses are provided in Supplemental Table 2.

### PAP administration did not impact the intraspinal inflammatory cell response after cervical SCI

Previous studies have shown that PTEN inhibition can modulate inflammatory responses following CNS injury (Liu et al., 2019; Pearson et al., 2023). Furthermore, intraspinal inflammation following SCI can impact outcomes such as secondary degeneration and axon growth capacity. Therefore, we performed immunohistochemistry (IHC) analysis for Iba1 to quantify the microglia/macrophage response at various gray matter locations at two caudal distances from the lesion epicenter (Figure 5A-G). Across the dorsal horn, intermediate, and ventral horn gray matter of the ipsilateral cervical spinal cord, Iba1 signal intensity was greater at the peri-lesional location than at the more distal/caudal intact location. However, at all examined locations, there was no difference between DMSO-only and PAP (Figure 5H). Statistical details for Iba1 analysis are provided in Supplemental Table 2. Together with the lesion and LMN analyses, these findings suggest that PAP’s ability to promote forelimb motor recovery was not mediated by reducing secondary degeneration or decreasing the inflammatory cell response.

**Figure 5.**
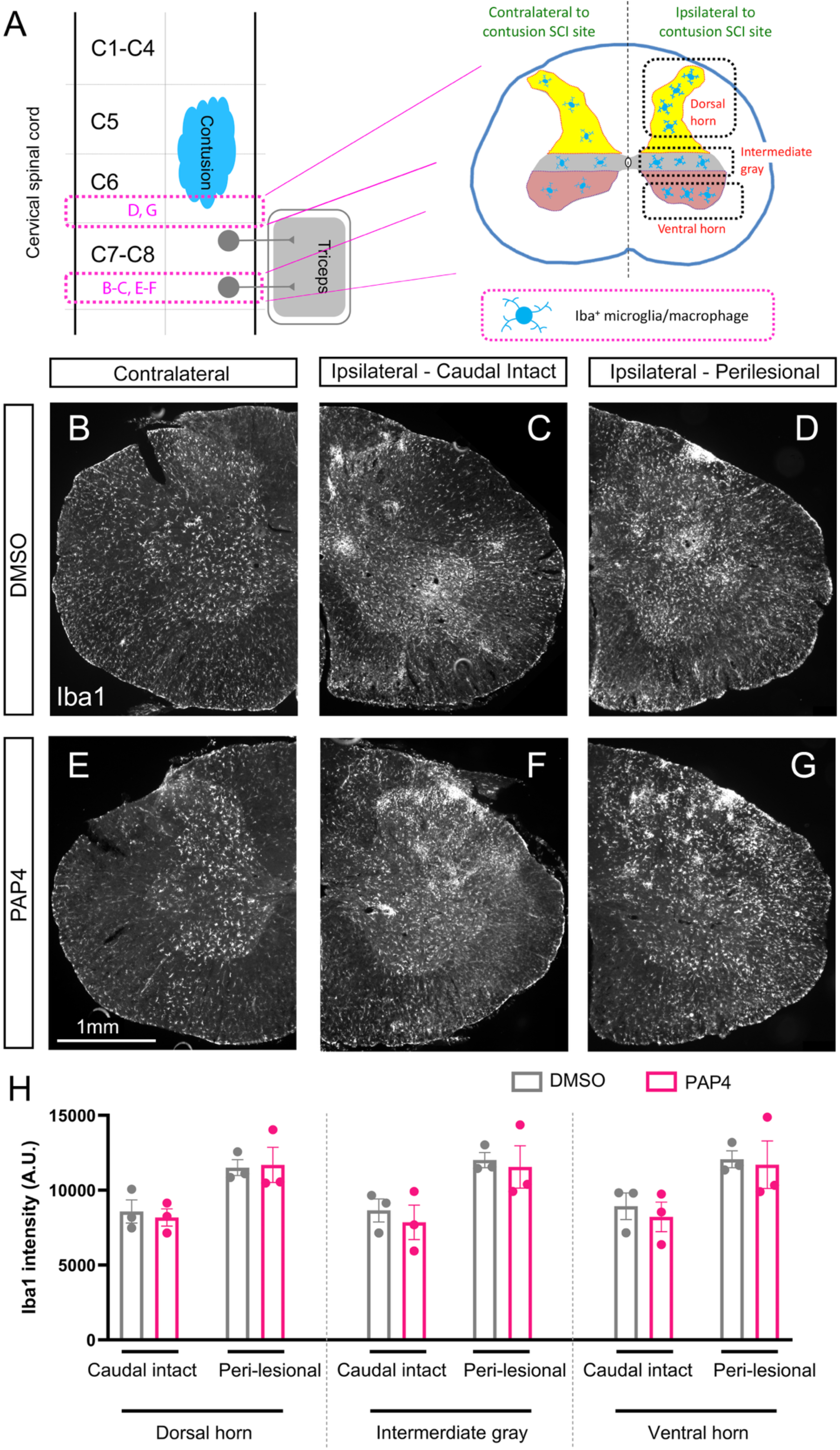
Iba1 analysis following chronic cervical contusion SCI and PAP treatment. Schematic depicting rostrocaudal and gray matter regions used for Iba1 fluorescence intensity analysis (A). Representative Iba1 immunofluorescence images from DMSO-only animals in the contralateral spinal cord (B), intact spinal cord caudal to the lesion (C), and perilesional region (D). Representative Iba1 immunofluorescence images from PAP-treated animals in the corresponding contralateral (E), caudal intact (F), and perilesional regions (G). Quantitative analysis of mean Iba1 fluorescence intensity in the dorsal horn, intermediate gray, and ventral horn within caudal intact and perilesional regions showed no significant differences between treatment groups (H). Details of statistical analyses are provided in Supplemental Table 2.

### PAP did not promote serotonergic axon growth following cervical contusion SCI

Given the role played by PTEN signaling in limiting axon growth capacity, we next examined the serotonergic axon growth response following cervical contusion SCI with PAP administration. Descending serotonergic axon projections provide important neuromodulatory input to motor circuitry of the cervical spinal cord and contribute to control of forelimb motor function. To assess whether the observed improvement in forelimb behavior was associated with increased serotonergic axon growth, we performed IHC for 5-HT (serotonin) and quantified density of 5-HT axons in the ipsilateral ventral horn caudal to the lesion site (Figure 6A), which is the location of the LMN pools innervating relevant forelimb muscles such as triceps. Both SCI groups showed a significant reduction in 5-HT axon density compared to uninjured animals; however, there was no difference between DMSO-only and PAP rats (Figure 6B-E). Statistical details for 5-HT axon analysis are provided in Supplemental Table 2. These findings show that PAP did not increase growth of 5-HT axons caudal/distal to the lesion.

**Figure 6.**
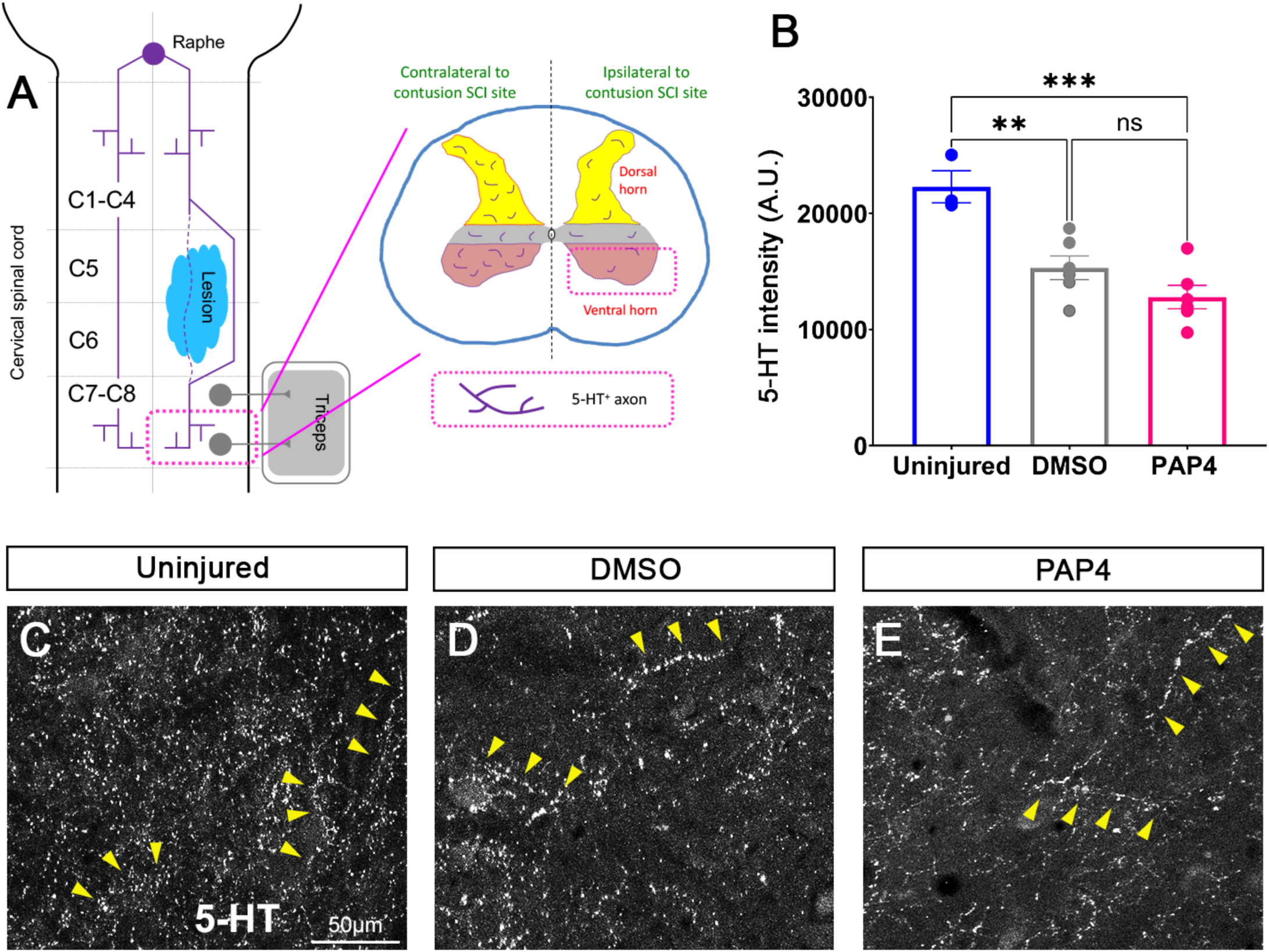
Serotonergic axon growth analysis following chronic cervical contusion SCI and PAP treatment. Schematic illustrating supraspinal 5-HT axon input and the caudal ventral horn region used for 5-HT fluorescence intensity analysis (**A).** Mean 5-HT fluorescence intensity in the ipsilateral caudal ventral horn was significantly higher in uninjured animals than in DMSO-only and PAP-treated animals, whereas the treatment groups did not differ significantly from one another **(B).** Representative 5-HT immunofluorescence images from ipsilateral caudal ventral horn of uninjured **(C),** DMSO-only **(D),** and PAP-treated **(E)** animals. ns, not significant (*p* > 0.05); \**p* < 0.05; \*\**p* < 0.001. Details of statistical analyses are provided in Supplemental Table 2.

### PAP promoted recruitment of multiple populations of intraspinal interneurons

Spinal cord interneurons play highly important roles in control of motor function. In the context of regenerative interventions after SCI, treatments may be able to stimulate functional recovery by targeting circuit plasticity mechanisms in addition to long-distance regeneration of the damaged axons of supraspinal neuron populations. Spinal interneurons are highly relevant to such plasticity, as they can serve as a substrate to generate polysynaptic relay circuitry (Bareyre et al., 2004). Specifically, supraspinal and/or other neuron populations could project their axons to innervate at-level projecting and intersegmental/propriospinal interneurons, which can then connect to the ventral horn LMNs that innervate forelimb muscle. At 22 weeks post-injury, animals were injected with the trans-synaptic retrograde tracer, pseudorabies virus (PRV-152; GFP reporter expressed in PRV-infected neurons), into triceps muscle ipsilateral to the SCI site to label forelimb-innervating LMNs in the cervical spinal cord. After first infecting these LMNs, the PRV-GFP then retrogradely transferred across the synapse to any neuron that made a synaptic connection with these LMNs. This approach allowed us to characterize polysynaptic relay circuitry that controls forelimb motor function.

Importantly, characterization of the organization of this triceps interneuron circuitry has not been conducted to date; therefore, we performed extensive quantitative PRV-GFP analysis of uninjured animals to first understand the circuitry in the intact nervous system condition. We found that a large portion of PRV-GFP labeled spinal interneurons that are part of triceps circuitry were located at the same spinal cord levels (C7-C8) as triceps-innervating LMNs, including in both ipsilateral and contralateral spinal cord (Figure 7A-B). While PRV-labeled locally projecting interneurons were broadly distributed across C7-C8 spinal gray matter, the largest number were in the dorsal horn compared to intermediate gray and ventral horn (Figure 7B). In addition, a greater proportion of these C7-C8 interneurons were located ipsilateral to PRV-injected muscle (Figure 7B). In the uninjured condition, PRV-positive interneurons were also found intersegmentally at levels far rostral (C2-C3) to the LMNs, both ipsilateral and contralateral to the muscle site of PRV-GFP injection (Figure 8A-B). Similar to the distribution in C7-C8, PRV-labeled intersegmentally projecting interneurons were identified across all gray matter locations at C2-C3, with the dorsal horn again being the site of the largest number, though they were present here at approximately half the numbers found at C7-C8 (Figure 8B). Together, these findings demonstrate that triceps interneuron circuitry includes both locally projecting and propriospinal populations distributed extensively throughout the cervical spinal cord.

**Figure 7.**
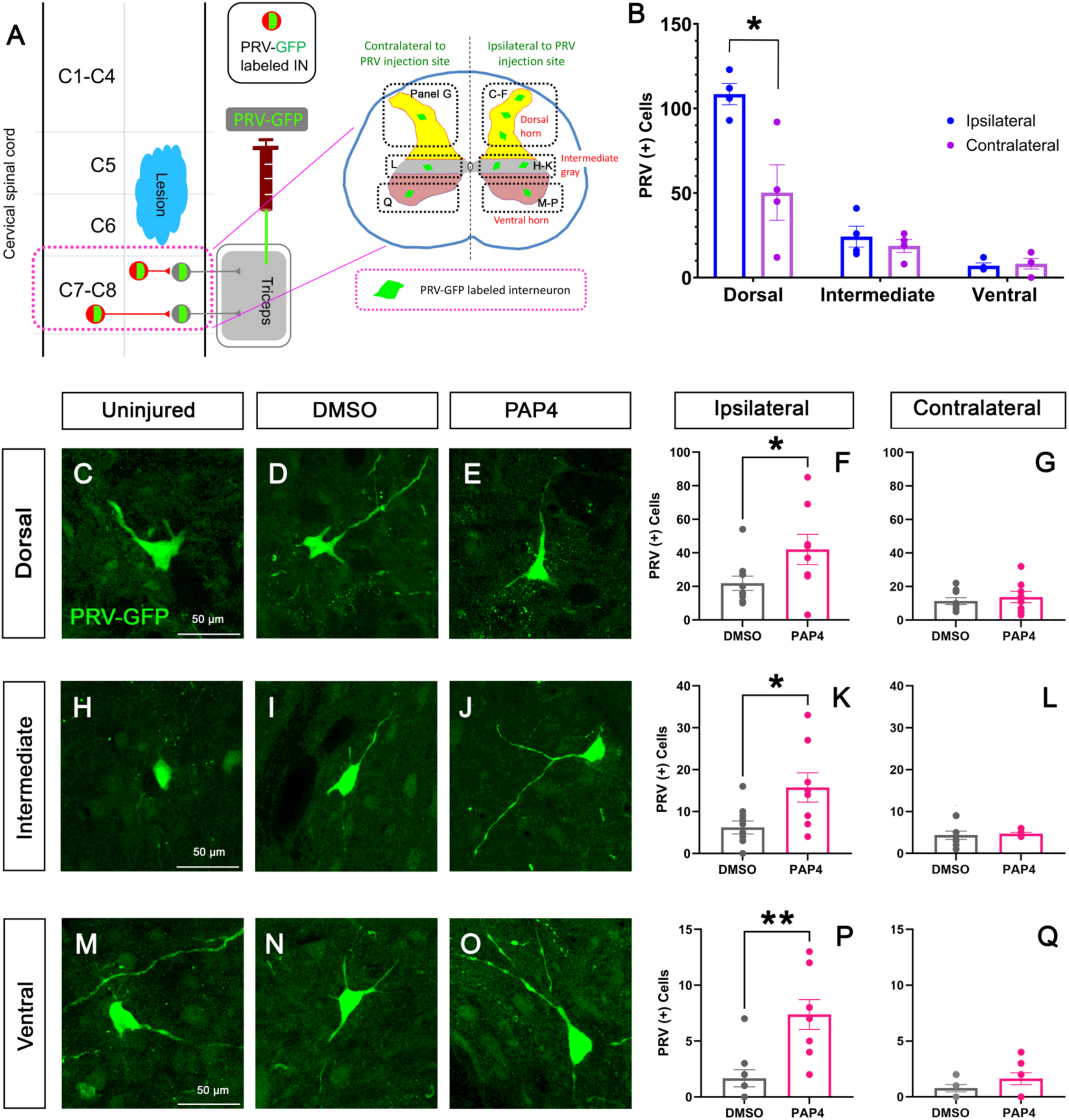
Caudal intraspinal interneuron circuitry following chronic cervical contusion SCI and PAP treatment. Schematic showing the location of PRV-152 injection and interneuron quantification caudal to the lesion within defined gray matter regions **(A).** Distribution of locally projecting spinal interneurons in uninjured animals on the ipsilateral and contralateral sides **(B).** Representative images of PRV-positive locally projecting spinal interneurons in the dorsal horn **(C–E),** intermediate gray **(H–J),** and ventral horn **(M–O)** across experimental groups. Quantification in the dorsal horn showed a significant increase in PRV-positive cell counts on the ipsilateral side with PAP, with no significant difference on the contralateral side **(F, G).** Quantification in the intermediate gray showed a significant increase on the ipsilateral side with PAP, with no significant difference on the contralateral side **(K, L).** Quantification in the ventral horn showed a significant increase on the ipsilateral side with PAP, with no significant difference on the contralateral side (P, Q). \**p* < 0.05. Details of statistical analyses are provided in Supplemental Table 3.

**Figure 8.**
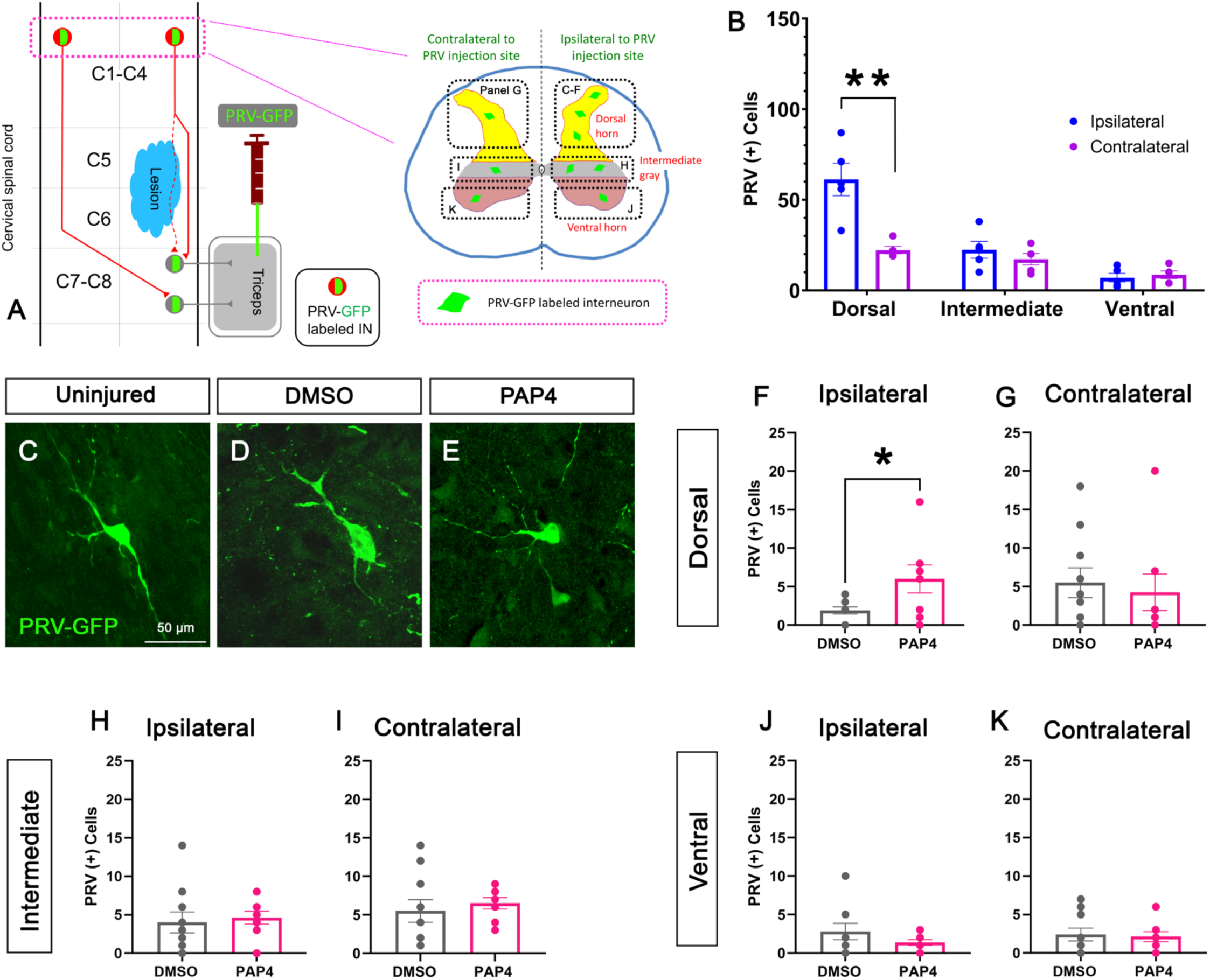
Propriospinal interneuron circuitry following chronic cervical contusion SCI and PAP treatment. Schematic showing the location of PRV-152 injection and propriospinal interneuron quantification rostral to the lesion within defined gray matter regions **(A).** Distribution of propriospinal interneurons in uninjured animals on the ipsilateral and contralateral sides **(B).** Representative images of PRV-positive propriospinal interneurons in the dorsal horn across experimental groups **(C-E).** Quantification in the dorsal horn showed a significant increase in PRV-positive cell counts on the ipsilateral side with PAP, with no significant difference on the contralateral side **(F, G).** Quantification in the intermediate gray **(H, I)** and the ventral horn **(J, K)** showed no significant differences on either the ipsilateral or contralateral side. \**p* < 0.05. Details of statistical analyses are provided in Supplemental Table 3.

We next asked whether PAP promoted plasticity of these spinal interneuron networks. Again, using quantitative analysis of PRV-GFP tracing, we found that cervical contusion SCI significantly reduced connectivity of spinal interneuron circuitry in both DMSO-only and PAP-treated groups. PAP promoted recruitment of both local and intersegmental projecting spinal interneuron circuits compared to DMSO-only control. Caudal to injury in C7-C8 (Figure 7A), PAP significantly increased PRV-GFP positive interneuron counts in ipsilateral dorsal horn (Figure 7D-F; *p* = 0.047), intermediate gray (Figure 7I-K; *p* = 0.017), and ventral horn (Figure 7N-P; *p* = 0.002), while we observed no differences between the two injury groups in corresponding contralateral gray matter regions (Figure 7G, L, Q). Statistical details for PRV analysis caudal to the lesion are provided in Supplemental Table 3. Rostral to injury (Figure 8A), PAP significantly increased numbers of PRV-GFP labeled long-distance projecting propriospinal interneurons present in ipsilateral C2-C3 dorsal horn (Figure 8D-F; *p* = 0.028), while we observed no significant differences between DMSO-only and PAP in intermediate gray (Figure 8H) or ventral horn (Figure 8J) of ipsilateral spinal cord or in corresponding contralateral C2-C3 gray matter regions (Figure 8G, I, K). Statistical details for PRV analysis rostral to the lesion are provided in Supplemental Table 3. Together, these PRV trans-synaptic tracing findings demonstrate that forelimb motor recovery after chronic cervical SCI promoted by PAP-based PTEN inhibition was associated with significantly enhanced connectivity of multiple spinal interneuron populations with forelimb-innervating LMN populations.

### PAP administration promoted formation of polysynaptic relay circuitry

As PAP enhanced connectivity of propriospinal interneurons located rostral to the lesion with triceps-innervating LMNs found caudal to the injury (Figure 8F), we examined synaptic input to these propriospinal interneurons. To do so, we performed IHC for PRV-GFP (to identify cell bodies of trans-synaptically labeled spinal interneurons), 5-HT (to label serotonergic axons, including all the way to their pre-synaptic terminals), and synaptophysin (a pan-synaptic marker to identify synaptic sites) (Figure 9A-E). In the ipsilateral C2-C3 dorsal horn, PAP significantly increased the number of synaptophysin-positive synaptic inputs to PRV-GFP-labeled propriospinal interneurons that were connected to triceps innervating C7-C8 LMNs compared to DMSO-only control (Figure 9F; *p* = 0.048). We also found that descending serotonergic axons made synaptic connections directly onto PRV-GFP labeled C2-C3 propriospinal interneurons, demonstrating a polysynaptic circuit: raphe to upper cervical propriospinal interneurons to lower cervical LMNs and ultimately to triceps. Compared to DMSO-only, PAP did not alter the number of 5-HT/synaptophysin double-positive serotonergic synaptic inputs per PRV-GFP labeled spinal interneuron in the ipsilesional C2-C3 dorsal horn (Figure 9G). As these analyses of synaptophysin and 5-HT/synaptophysin synapses were performed on a per PRV-GFP-labeled interneuron basis, these data do not reflect the even greater increase in overall connectivity of this polysynaptic circuitry due to the enhanced connection between C2-C3 propriospinal interneurons and C7-C8 LMNs (as evidenced in Figure 8F by the greater number of total PRV-GFP labeled interneurons rostral to the injury with PAP). Statistical details for synaptic analysis are provided in Supplemental Table 3. Together, these results suggest that PAP promoted recruitment of polysynaptic intersegmental relay circuitry that rostro-caudally spanned the SCI site over multiple segments.

**Figure 9.**
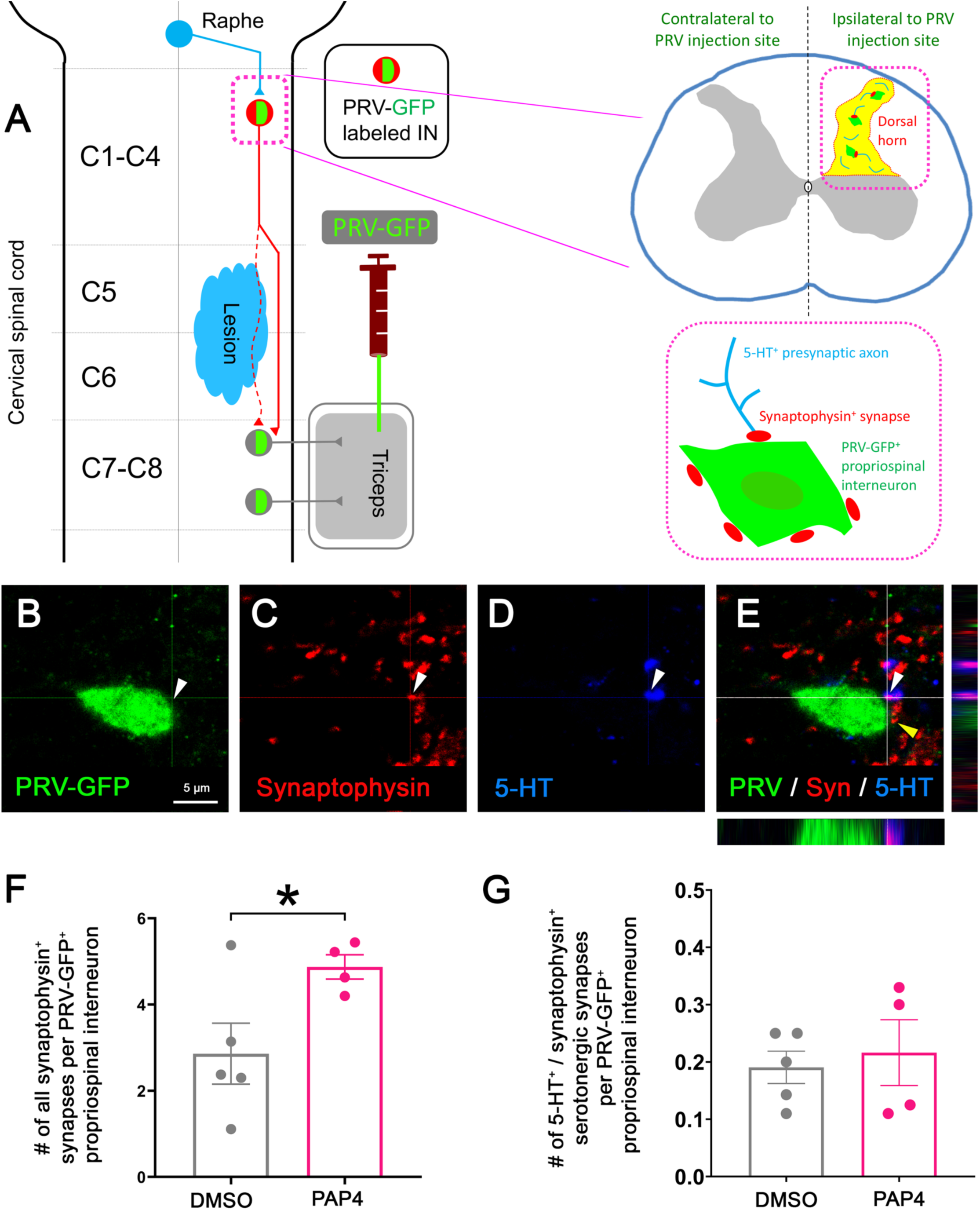
Synaptic remodeling within forelimb interneuron circuitry following chronic cervical contusion SCI and PAP treatment. Schematic illustrating supraspinal 5-HT axon input onto PRV-labeled forelimb-associated propriospinal interneurons in the rostral dorsal horn region used for analysis **(A).** Representative confocal images showing PRV-GFP **(B),** synaptophysin **(C),** 5-HT **(D),** and merged labeling **(E).** Quantification of synaptophysin-positive puncta associated with each PRV-positive propriospinal interneuron cell body was significantly increased following PAP treatment **(F).** Quantification of 5-HT^+^/synaptophysin^+^ puncta associated with each PRV-positive cell body showed no significant difference between treatment groups **(G).** \**p* < 0.05. Details of statistical analyses are provided in Supplemental Table 3.

### Effect of cervical SCI and PAP on supraspinal connectome of triceps motor circuitry

While a major focus in the SCI field has been placed on CST plasticity when attempting to promote motor recovery, a number of neuron populations in the brain that project descending axons to the spinal cord operate together as a group to control motor function, including for the forelimb (Blackmore et al., 2021). To determine whether PAP administration altered connectivity of these supraspinal neuron populations with triceps motor circuitry located caudal to the SCI site, we again utilized PRV trans-synaptic tracing (Figure 10A). As characterization of the supraspinal “connectome” for triceps motor circuitry using PRV tracing has not previously been conducted, we first performed extensive quantitative PRV analysis to determine which neuron populations of the brain are connected to triceps motor circuitry in uninjured animals. We found PRV-labeled neurons in a number of nuclei across the brainstem (Figure 10B-L). The largest number of PRV-labeled brainstem neurons were found in the medullary reticular formation, with prominent labeling present in the more rostrally located gigantocellular reticular nucleus (Gi; Figure 10E-F), gigantocellular reticular nucleus alpha and ventral parts (GiA and GiV; Figure 10E,G), and lateral paragigantocellular nucleus alpha and lateral parts (LPGiA and LPGi; Figure 10 E,H). PRV-labeled neurons were also identified more caudally within the dorsal and ventral medullary reticular nuclei (MdD/MdV; Figure 10J-K). We additionally found substantial PRV labeling within the pontine reticular formation, including the caudal pontine reticular nucleus (PnC; Figure 10B,D) and ventral pontine reticular nucleus (PnV; Figure 10B-C). In each of these bilaterally organized nuclei, we observed similar levels of PRV labeling on both sides of the brainstem in uninjured animals (Figure 10L). In addition, PRV-labeled neurons were identified within midline localized nuclei of the caudal raphe, including raphe obscurus (ROb), raphe pallidus (RPa), and raphe magnus (RMg) (Figure 10E,I,L).

**Figure 10.**
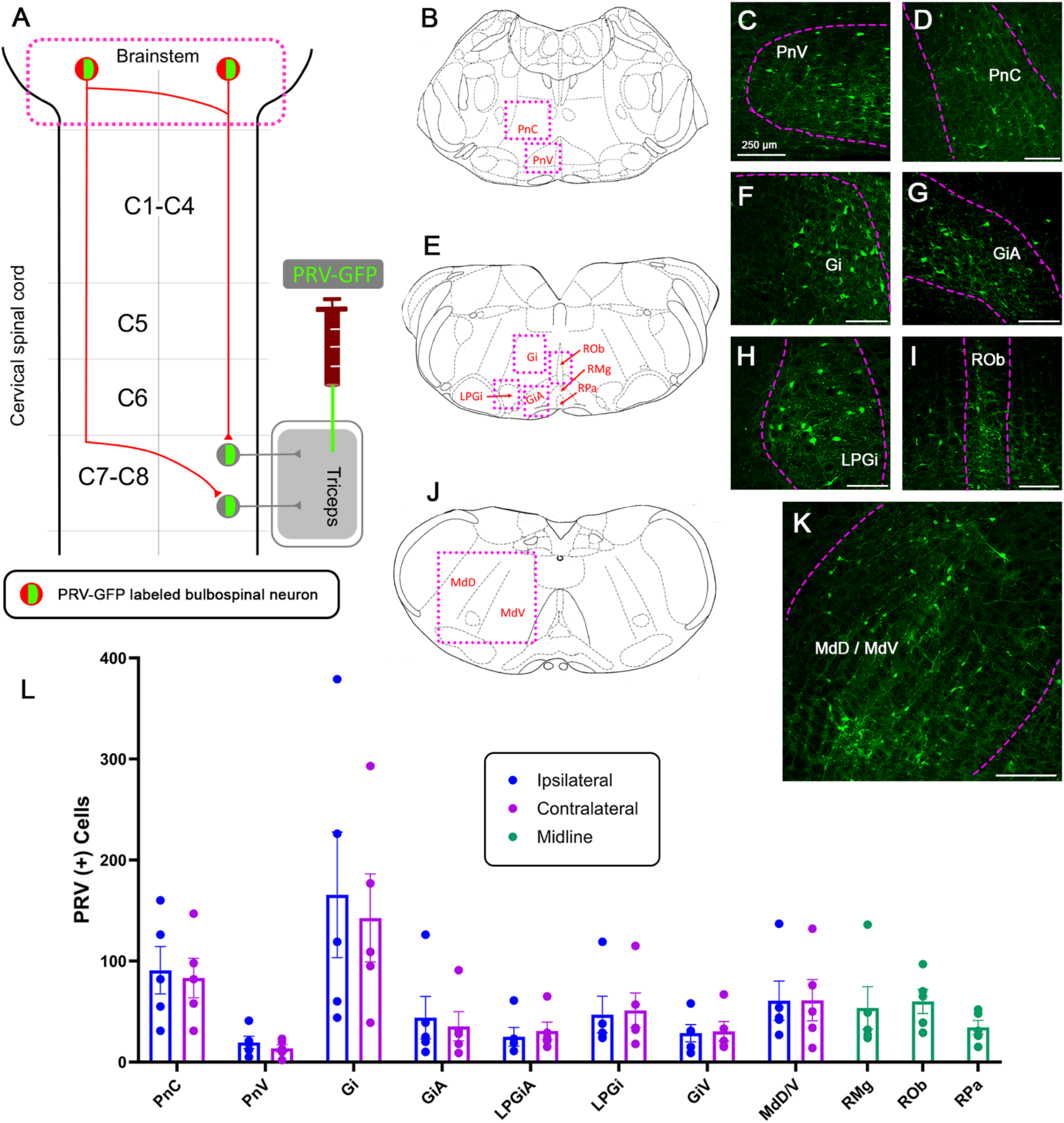
Supraspinal brainstem populations connected with triceps motor circuitry in uninjured animals. Schematic illustrating PRV-152 injection into the triceps muscle and retrograde trans-synaptic labeling of supraspinal neuron populations associated with forelimb motor circuitry **(A).** Representative brainstem schematics identifying analyzed regions within the pontine reticular formation **(B),** rostral medulla **(E),** and caudal medulla **(J)**; modified from https://labs.gaidi.ca/rat-brain-atlas. Representative PRV-GFP labeling in the caudal and ventral pontine reticular nuclei (PnC/PnV) **(C**-**D),** gigantocellular reticular nucleus (Gi) **(F),** gigantocellular reticular nucleus alpha (GiA) **(G),** lateral paragigantocellular nucleus (LPGi) **(H),** raphe obscurus (ROb) **(I),** and dorsal and ventral medullary reticular nuclei (MdD/V) **(K).** Distribution of PRV-positive supraspinal neurons associated with forelimb motor circuitry across ipsilateral, contralateral, and midline brainstem regions in uninjured animals **(L).** Details of statistical analyses are provided in Supplemental Table 4.

Following cervical contusion SCI, numbers of PRV-labeled neurons were broadly reduced across all examined brainstem nuclei relative to levels observed in uninjured animals, including on both ipsilateral and contralateral sides of the brainstem (Figure 11). We next compared PRV-positive neuron numbers between DMSO- and PAP-treated SCI animals to determine whether PAP altered recruitment of these supraspinal populations into forelimb-associated circuitry. No significant differences between DMSO and PAP were detected within PnC (Figure 11A), PnV (Figure 11B), Gi (Figure 11C), GiA (Figure 11D), LPGiA (Figure 11E), LPGi (Figure 11F), GiV (Figure 11G), or MdD/MdV (Figure 11H) on either the ipsilateral or contralateral side. Similarly, PAP treatment did not alter PRV-positive neurons numbers within the midline raphe nuclei RMg (Figure 11I), ROb (Figure 11J), or RPa (Figure 11K). Statistical details for brainstem PRV analysis are provided in Supplemental Table 4.

**Figure 11.**
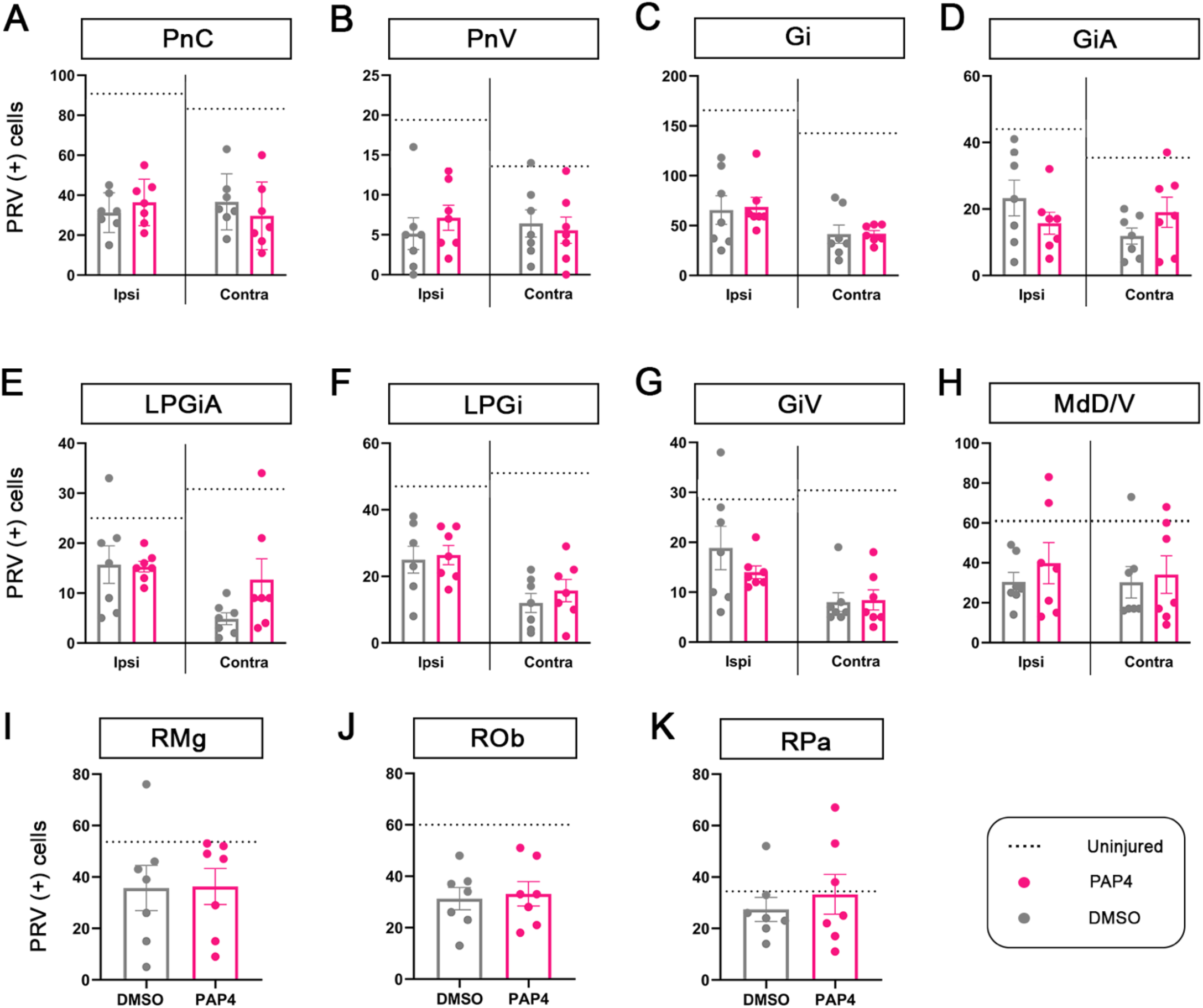
Supraspinal brainstem populations connected with triceps motor circuitry in chronic cervical contusion SCI and in response to PAP treatment. Quantification of PRV-positive neurons within the caudal pontine reticular nucleus (PnC) **(A),** ventral pontine reticular nucleus (PnV) **(B),** gigantocellular reticular nucleus (Gi) **(C),** gigantocellular reticular nucleus alpha (GiA) **(D),** lateral paragigantocellular nucleus alpha (LPGiA) **(E),** lateral paragigantocellular nucleus (LPGi) **(F),** ventral gigantocellular reticular nucleus (GiV) **(G),** and dorsal and ventral medullary reticular nuclei (MdD/V) **(H)** on the ipsilateral and contralateral sides. Quantification of PRV-positive neurons within the midline raphe magnus (RMg) (**I),** raphe obscurus (ROb) **(J),** and raphe pallidus (RPa) **(K).** Dashed lines indicate the corresponding mean PRV-positive cell count observed in uninjured animals and illustrate the overall reduction in supraspinal labeling following cervical SCI. No significant differences in PRV-positive cell counts were detected between DMSO-only and PAP-treated animals in any analyzed brainstem nucleus. Details of statistical analyses are provided in Supplemental Table 4.

Together, these findings with PRV trans-synaptic tracing demonstrate that a large number of supraspinal neuron populations contribute to the descending circuitry associated with triceps motor output, showing the high degree of supraspinal connectome complexity controlling forelimb motor function. Furthermore, cervical SCI was associated with a broad reduction in PRV labeling across these brainstem populations at a chronic time point, and PAP did not significantly alter PRV labeling within these supraspinal populations.

**Figure 12.**
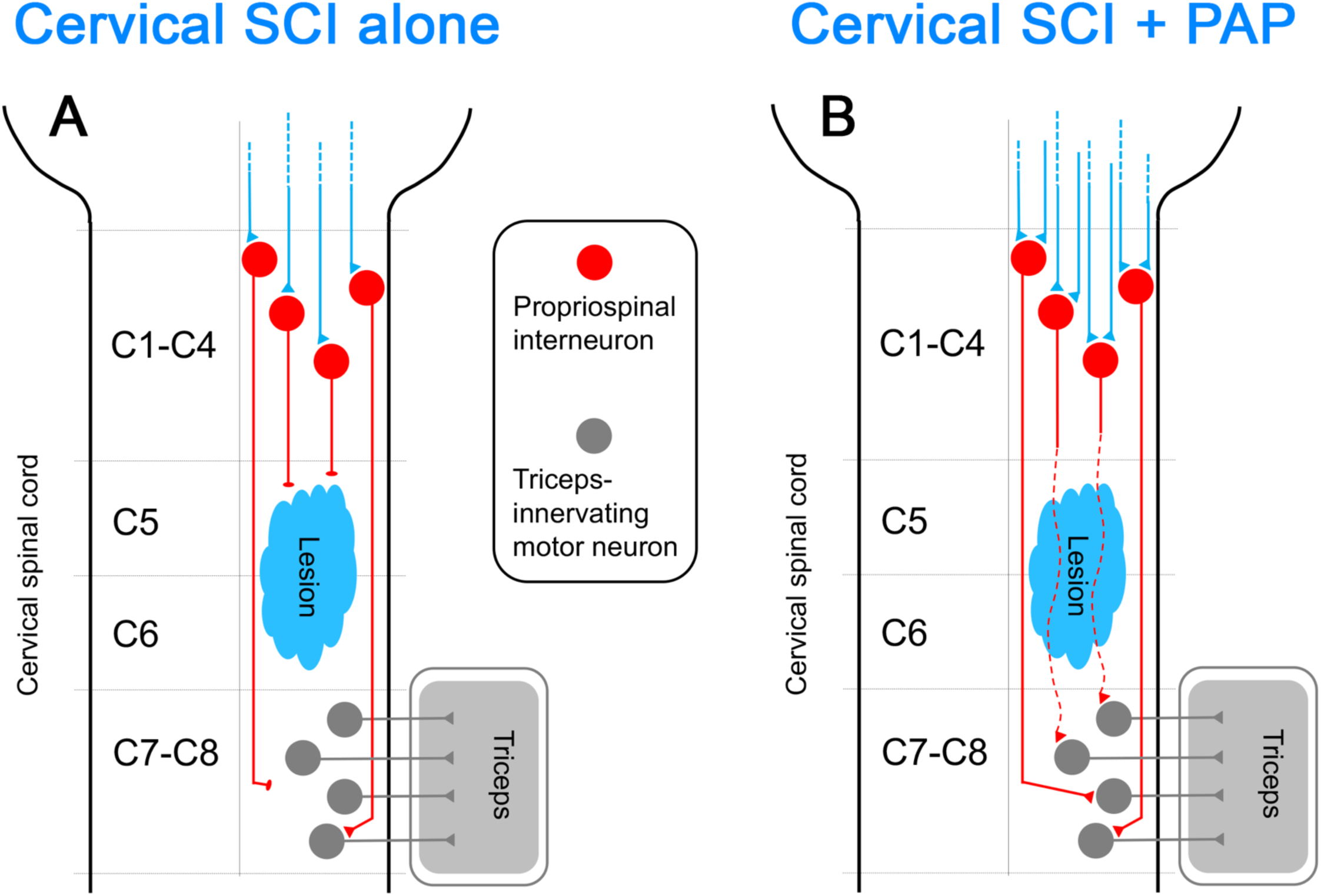
Summary diagram. Following cervical contusion SCI, there is significant disruption of spinal interneuron connectivity within forelimb motor circuitry, including intersegmentally projecting propriospinal populations. In response to PAP treatment at a chronic point post-injury, these interneuron populations are recruited to enhance their synaptic connectivity with forelimb-innervating lower motor neurons located caudal to the SCI site. This recruitment may allow for generation of polysynaptic circuitry to relay descending input from supraspinal neuron populations that are important for control of forelimb motor function.

## DISCUSSION

In this study, we investigated whether pharmacologic PTEN inhibition could induce functional motor recovery and plasticity of forelimb motor circuitry after chronic cervical contusion SCI. We found that PAP promoted selective recovery in only a subset of functionalities, with improvements in forelimb usage during cylinder testing that were not observed across other behavioral assays. This recovery occurred in the absence of detectable differences in lesion size, LMN survival, inflammatory response, or serotonergic axon innervation caudal to the injury. In contrast, PAP resulted in recruitment of multiple spinal interneuron populations, including both local-projecting and propriospinal neurons, within forelimb motor circuitry. We also observed an increase in synaptic input onto these recruited propriospinal interneurons. In addition, this study characterized a broad supraspinal network associated with forelimb motor circuitry, including pontine reticular, medullary reticular, and raphe nuclei. These supraspinal connectome findings further define the distributed organization of forelimb-associated motor circuitry. Together, our results demonstrate that remodeling of spinal interneuron networks may be an important circuit plasticity mechanism to target for promoting forelimb motor recovery in chronic cervical SCI.

PAP administration promoted selective recovery of forelimb usage during rearing events, as assessed by cylinder paw-preference testing, but not in measures of forelimb grip strength, sensorimotor function, or gait-related parameters. Cylinder testing is generally considered a measure of gross forelimb usage rather than skilled motor performance, suggesting that the behavioral benefit of PAP may have been due to preferential impact on circuitry supporting gross and postural motor output (Fouad et al., 2020). These functions are strongly influenced by bulbospinal systems, including reticulospinal, vestibulospinal, and serotonergic pathways, which provide major descending input to spinal motor networks (Fouad et al., 2020; Akalu et al., 2023; Giorgi et al., 2025). Interestingly, Filli et al. (2014) demonstrated that injured reticulospinal fibers can sprout onto cervical propriospinal neurons that bypass the lesion and reconnect with denervated spinal circuitry, forming a “reticulo-propriospinal detour” associated with functional motor recovery after cervical SCI. Consistent with this model, our data with therapeutic manipulation in a chronic contusion paradigm suggest that PAP-induced recovery may reflect remodeling of the spinal interneuron circuits through which these descending bulbospinal systems act.

In addition to increased connectivity between cervical interneuron networks and forelimb-innervating LMNs, we also found that PAP increased synaptic input to recruited propriospinal interneurons. This raises the possibility that the observed recovery of cylinder performance was supported by promoting formation of spinal relay circuits. While we found serotonergic synaptic input to PRV-labeled propriospinal interneurons, potential additional sources of presynaptic input include other bulbospinal systems, such as reticulospinal projections, which are known to connect with propriospinal circuitry after cervical SCI (Filli et al., 2014). In addition, because PAP increased the overall number of recruited interneurons, the total serotonergic influence across this circuitry could still be enhanced even if the serotonergic synaptic input to each individual interneuron was unchanged. This possibility would be in line with previous work showing that serotonergic modulation can facilitate forelimb motor recovery after cervical SCI (Jin et al., 2021). Together, these findings suggest that PAP may support recovery by increasing both the recruitment of spinal interneurons and amount of descending synaptic input they receive.

While CST plasticity has been extensively investigated in the recovery of forelimb motor function after cervical SCI, the contribution of bulbospinal pathways remains highly understudied (Bareyre et al., 2004; Anderson et al., 2009; Du et al., 2015). Wang and colleagues (2018; 2022) demonstrated that cervical spinal cord receives input from a widely distributed network of supraspinal populations, including multiple reticulospinal and vestibulospinal nuclei, emphasizing that forelimb motor control extends well beyond CST input. In the present study, we provide an in-depth characterization of supraspinal brainstem populations associated with triceps motor circuitry. PRV labeling was distributed across both ipsilateral and contralateral brainstem populations and was reduced bilaterally following cervical SCI, highlighting the complex organization of supraspinal input to forelimb motor networks. Together with the increased recruitment and synaptic input observed within spinal interneuron populations, these findings further support the idea that PAP-induced plasticity may have occurred primarily through remodeling of spinal relay networks rather than increased direct connectivity of supraspinal neurons with LMNs.

Previous work has shown that combining PTEN inhibition with task-based rehabilitative training can further enhance circuit remodeling and forelimb motor recovery after cervical SCI compared to PTEN inhibition alone (Pan et al., 2021). This suggests that enhancing neuron-intrinsic growth capacity may prime injured and spared circuits for plasticity, and rehabilitative training may translate this increased plasticity into greater functional recovery. Future studies could therefore determine whether combining PAP treatment with rehabilitative training produces broader circuit reorganization and functional recovery in our model. Additionally, future work will be necessary to identify the supraspinal sources of the increased synaptic input onto PAP-recruited spinal interneurons and to determine whether these pathways are required for functional recovery. One approach would be to anterogradely label candidate bulbospinal populations and determine whether PAP enhances the synaptic connection between these axon projections and PRV-labeled spinal interneurons. Functional manipulation of these pathways, including selective silencing, could further determine whether bulbospinal input to spinal relay circuitry is necessary for the behavioral recovery observed in this study.

We conducted this study in the chronic phase of cervical SCI. Most regenerative strategies following SCI have been investigated during the acute or subacute periods, despite the vast majority of individuals living with chronic injury (Tashiro et al., 2017). Promoting plasticity at chronic time points can be particularly challenging because of persistent neuron-extrinsic and -intrinsic growth inhibition (Tran et al., 2018). Despite these barriers, our findings emphasize that enhancing axon growth capacity at chronic time points after injury using a non-invasive pharmacological approach can still promote functional motor recovery and significant remodeling of spared circuitry. This is particularly relevant following cervical SCI, where recovery of upper extremities is a major priority for those with chronic injury (Anderson, 2004). Together, these findings support the continued development of strategies that target plasticity of spinal motor circuits to promote recovery well into the chronic phase of SCI.

## ACKNOWLEDGMENTS

This study was funded by the NINDS (R01NS079702 and R01NS110385 to A.C.L.) and the Yant Family Spinal Cord Regeneration Fund. We thank the Princeton Neuroscience Institute Viral Core Facility for providing PRV. The authors would especially like to acknowledge Mr. Bob Yant for his encouragement, generous support of this work, and passion for finding therapies to promote functional recovery after chronic spinal cord injury.

## AUTHOR CONTRIBUTIONS (CRediT)

Conceptualization: K Crowther, AC Lepore

Data curation: K Crowther, AC Lepore

Formal analysis: K Crowther, A Harbeck, M Middlesworth, AC Lepore

Funding acquisition: AC Lepore

Investigation: K Crowther, A Harbeck, M Middlesworth, M Bosch, MA Lyttle, C MacGibbon, DA Jaffe, P Michel-Flutot, E Fitchett, AC Lepore

Methodology: K Crowther, A Harbeck, M Middlesworth, M Bosch, E Fitchett, AC Lepore Project administration: AC Lepore

Resources: S Li, AC Lepore Software: N/A

Supervision: AC Lepore

Validation: K Crowther, A Harbeck, M Middlesworth, AC Lepore

Visualization: K Crowther, A Harbeck, M Middlesworth, AC Lepore

Roles/Writing – original draft: K Crowther, AC Lepore

Writing – review & editing: K Crowther, AC Lepore

## ABBREVIATIONS

5-HT: 5-hydroxytryptamine (serotonin)
CST: corticospinal tract
C2, C3, etc.: cervical spinal cord level 2, 3, etc.
DMSO: dimethyl sulfoxide
GFP: green fluorescent protein
Gi: gigantocellular reticular nucleus
GiA: gigantocellular reticular nucleus, alpha part
GiV: gigantocellular reticular nucleus, ventral part
Iba1: ionized calcium-binding adapter molecule 1
IHC: immunohistochemistry
LMN: lower motor neuron
LPGi: lateral paragigantocellular reticular nucleus
LPGiA: lateral paragigantocellular reticular nucleus, alpha part
MdD: medullary reticular nuclei, dorsal part
MdV: medullary reticular nuclei, ventral part
PAP: PTEN antagonist peptide
PnC: caudal pontine reticular nucleus
PnV: ventral pontine reticular nucleus
PRV: pseudorabies virus
PTEN: phosphatase and tensin homolog
RMg: raphe magnus
ROb: raphe obscurus
RPa: raphe pallidus
SCI: spinal cord injury

---

**Supplemental table 1.**

| Figure panel | Outcome | Analysis | Comparison | N1, N2 | Mean 1 ± SEM | Mean 2 ± SEM | Test statistic | P value |
|---|---|---|---|---|---|---|---|---|
| Fig. 2B | Ipsilateral grip strength | Mixed-effects model | Treatment | — | — | — | F (1, 28) = 0.9338 | 0.3422 |
|  |  |  | Time | — | — | — | F (3.171, 69.77) = 11.99 | **<0.0001** |
|  |  |  | Treatment x Time | — | — | — | F (4, 88) = 0.9915 | 0.4165 |
| Fig. 2C | Contralateral grip strength | Mixed-effects model | Treatment | — | — | — | F (1, 28) = 0.91 | 0.3483 |
|  |  |  | Time | — | — | — | F (2.477, 57.59) = 2.785 | 0.0588 |
|  |  |  | Treatment x Time | — | — | — | F (4, 93) = 0.9319 | 0.4490 |
| Fig. 2E | Ipsilateral foot faults | Mixed-effects model | Treatment | — | — | — | F (1, 28)= 0.000009313 | 0.9976 |
|  |  |  | Time | — | — | — | F (1.922, 32.68) = 21.72 | **<0.0001** |
|  |  |  | Treatment x Time | — | — | — | F (2, 34) = 0.01393 | 0.9862 |
|  |  | Šidák's multiple comparisons | DMSO: Baseline vs 8wpi | 12, 11 | 8.5 ± 1.125 | 27 ± 4.171 | t (7) = 3.394 | **0.0342** |
|  |  |  | DMSO: Baseline vs 12wpi | 12, 10 | 8.5 ± 1.125 | 23.7 ± 3.229 | t (9) = 4.657 | **0.0036** |
|  |  |  | PAP4: Baseline vs 8wpi | 13, 12 | 9.3 ± 2.092 | 26.5 ± 2.770 | t (9) = 4.301 | **0.0060** |
|  |  |  | PAP4: Baseline vs 12wpi | 13, 10 | 9.3 ± 2.092 | 23.7 ± 4.803 | t (9) = 2.267 | 0.1417 |
| Fig. 2F | Contralateral Ffot faults | Mixed-effects model | Treatment | — | — | — | F (1, 28) = 0.7488 | 0.3942 |
|  |  |  | Time | — | — | — | F (1.919, 32.62) = 6.161 | **0.0059** |
|  |  |  | Treatment x Time | — | — | — | F (2, 34) = 0.2101 | 0.8116 |
|  |  | Šidák's multiple comparisons | DMSO: Baseline vs 8wpi | 12, 11 | 6.417 ± 1.203 | 13 ± 2.812 | t (7) = 1.604 | 0.3916 |
|  |  |  | DMSO: Baseline vs 12wpi | 12, 10 | 6.417 ± 1.203 | 9.2 ± 2.703 | t (9) = 0.8847 | 0.7833 |
|  |  |  | PAP4: Baseline vs 8wpi | 13, 12 | 4.692 ± 0.970 | 10.5 ± 1.782 | t (9) = 2.906 | 0.0513 |
|  |  |  | PAP4: Baseline vs 12wpi | 13, 10 | 4.692 ± 0.970 | 9.4 ± 1.529 | t (9) = 2.017 | 0.2073 |
| Fig. 2I | Stride Llngth | Mixed-effects model | Treatment | — | — | — | F (1, 28) = 0.2352 | 0.6315 |
|  |  |  | Time | — | — | — | F (1.772, 37.21) = 22.2 | **<0.0001** |
|  |  |  | Treatment x Time | — | — | — | F (1.772, 37.21) = 0.7013 | 0.4859 |
|  |  | Šidák's multiple comparisons | DMSO: Baseline vs 2wpi | 14, 11 | 137.976 ± 3.543 | 150.758 ± 4.318 | t (10) = 2.180 | 0.1542 |
|  |  |  | DMSO: Baseline vs 15wpi | 14, 11 | 137.976 ± 3.543 | 165.242 ± 4.648 | t (10) = 3.960 | **0.0080** |
|  |  |  | PAP4: Baseline vs 2wpi | 16, 13 | 138.833 ± 3.856 | 143.179 ± 5.531 | t (12) = 0.6765 | 0.8834 |
|  |  |  | PAP4: Baseline vs 15wpi | 16, 11 | 138.833 ± 3.856 | 166.303 ± 3.948 | t (10) = 4.641 | **0.0028** |
| Fig. 2J | Toe spread | Mixed-effects model | Treatment | — | — | — | F (1, 28) = 0.09548 | 0.7596 |
|  |  |  | Time | — | — | — | F (1.847, 39.70) = 13.98 | **<0.0001** |
|  |  |  | Treatment x Time | — | — | — | F (1.847, 39.70) = 3.419 | **0.0462** |
|  |  | Šidák's multiple comparisons | DMSO: Baseline vs 2wpi | 14, 13 | 18.576 ± 0.320 | 16.190 ± 0.543 | t (12) = 4.691 | **0.0016** |
|  |  |  | DMSO: Baseline vs 15wpi | 14, 11 | 18.576 ± 0.320 | 18.542 ± 0.378 | t (10) = 0.3778 | 0.9765 |
|  |  |  | PAP4: Baseline vs 2wpi | 16, 12 | 18.038 ± 0.295 | 17.319 ± 0.633 | t (11) = 1.078 | 0.6630 |
|  |  |  | PAP4: Baseline vs 15wpi | 16, 11 | 18.038 ± 0.295 | 18.412 ± 0.737 | t (10) = 0.8774 | 0.7849 |
| Fig. 2K | Surface area | Mixed-effects model | Treatment | — | — | — | F (1, 28) = 0.06951 | 0.7940 |
|  |  |  | Time | — | — | — | F (1.986, 46.67) = 10.03 | **0.0002** |
|  |  |  | Treatment x Time | — | — | — | F (1.986, 46.67) = 1.161 | 0.3217 |
|  |  | Šidák's multiple comparisons | DMSO: Baseline vs 2wpi | 14, 13 | 67.221 ± 3.160 | 59.801 ± 3.502 | t (12) = 1.367 | 0.4816 |
|  |  |  | DMSO: Baseline vs 15wpi | 14, 11 | 67.221 ± 3.160 | 70.446 ± 3.317 | t (10) = 1.054 | 0.6807 |
|  |  |  | PAP4: Baseline vs 2wpi | 16, 14 | 71.427 ± 3.492 | 54.602 ± 3.090 | t (13) = 3.271 | **0.0181** |
|  |  |  | PAP4: Baseline vs 15wpi | 16, 13 | 71.427 ± 3.492 | 73.559 ±5.498 | t (12) = 0.5719 | 0.9248 |
| Fig. 2M | Bilateral paw touches | Mixed-effects model | Treatment | — | — | — | F (1, 25) = 5.860 | **0.0231** |
|  |  |  | Time | — | — | — | F (2.470, 41.17) = 9.559 | **0.0002** |
|  |  |  | Treatment x Time | — | — | — | F (3, 50) = 3.747 | **0.0166** |
|  |  | Šidák's multiple comparisons | DMSO vs PAP4: Baseline | 14, 12 | 68.14 ± 3.856 | 63.08 ± 6.326 | t (18.53) = 0.6829 | 0.9390 |
|  |  |  | DMSO vs PAP4: 2wpi |  | 31.08 ± 7.406 | 35.17 ± 9.039 | t (21.18) = 0.3494 | 0.9947 |
|  |  |  | DMSO vs PAP4: 15wpi |  | 25.79 ± 2.076 | 59.94 ± 9.253 | t (8.796) = 3.601 | **0.0236** |
|  |  |  | DMSO vs PAP4: 18wpi |  | 27.50 ± 10.014 | 59.86 ± 8.105 | t (15) = 2.512 | 0.0924 |

**Supplemental table 2.**

| Figure panel | Outcome | Analysis | Comparison | N1, N2 | Mean 1 ± SEM | Mean 2 ± SEM | Test statistic | P value |
|---|---|---|---|---|---|---|---|---|
| Fig. 3H | Lesion area | Mixed-effects model | Treatment | — | — | — | F (1, 10) = 0.1394 | 0.7167 |
|  |  |  | Distance | — | — | — | F (3.393, 30.30) = 11.00 | **<0.0001** |
|  |  |  | Treatment x Distance | — | — | — | F (3.393, 30.30) = 0.3856 | 0.7874 |
| Fig. 3I | Lesion volume | Unpaired t-test | DMSO vs PAP4 | 6, 6 | 7.45 ± 0.71 | 7.125 ± 0.99 | t (10) = 0.2663 | 0.7955 |
| Fig. 4D | Lower motor neuron counts | Mixed-effects model | Treatment | — | — | — | F (1, 10) = 0.8509 | 0.3780 |
|  |  |  | Distance | — | — | — | F (3.716, 34.24) = 34.19 | **<0.0001** |
|  |  |  | Treatment x Distance | — | — | — | F (3.716, 34.24) = 0.8232 | 0.5121 |
| Fig. 5H | Iba1 fluorescence intensity | Unpaired t-test | Dorsal horn caudal intact:<br>DMSO vs PAP4 | 3, 3 | 8582 ± 771.7 | 8182 ± 577.0 | t (4) = 0.4148 | 0.6996 |
|  |  |  | Dorsal horn perilesional:<br>DMSO vs PAP4 | 3, 3 | 11511 ± 532.5 | 11693 ± 1174 | t (4) = 0.1409 | 0.8948 |
|  |  |  | Intermediate gray caudal intact:<br>DMSO vs PAP4 | 3, 3 | 8650 ± 766.7 | 7853 ± 1152 | t (4) = 0.5758 | 0.5956 |
|  |  |  | Intermediate gray perilesional:<br>DMSO vs PAP4 | 3, 3 | 12014 ± 509.7 | 11558 ± 1411 | t (4) = 0.3042 | 0.7761 |
|  |  |  | Ventral horn caudal intact:<br>DMSO vs PAP4 | 3, 3 | 8934 ± 886.7 | 8221 ± 985.9 | t (4) = 0.5382 | 0.6190 |
|  |  |  | Ventral horn perilesional:<br>DMSO vs PAP4 | 3, 3 | 12068 ± 567.6 | 11707 ± 1591 | t (4) = 0.2137 | 0.8412 |
| Fig. 6B | 5-HT fluorescence intensity | One-way ANOVA | Overall group effect | — | — | — | F (2, 12) = 14.8 | **0.0006** |
|  |  | Tukey's multiple comparisons | Uninjured vs DMSO | 3, 6 | 22289 ± 1383 | 15315 ± 1029 | q (12) = 5.631 | **0.0048** |
|  |  |  | Uninjured vs PAP4 | 3, 6 | 22289 ± 1383 | 12798 ± 1006 | q (12) = 7.663 | **0.0004** |
|  |  |  | DMSO vs PAP4 | 6, 6 | 15315 ± 1029 | 12798 ± 1006 | q (12) = 2.489 | 0.2242 |

**Supplemental table 3.**

| Figure panel | Outcome | Analysis | Comparison | N1, N2 | Mean 1 ± SEM | Mean 2 ± SEM | Test statistic | P value |
|---|---|---|---|---|---|---|---|---|
| Fig. 7B | Caudal PRV-positive cell counts | Unpaired t-test | Uninjured dorsal horn:<br>Ipsi vs Contra | 4, 4 | 108.5 ± 6.252 | 50.25 ± 16.42 | t (6) = 3.315 | **0.0161** |
|  |  |  | Uninjured intermediate gray:<br>Ipsi vs Contra | 4, 4 | 24.25 ± 6.169 | 18.75 ± 3.860 | t (6) = 0.7558 | 0.4784 |
|  |  | Mann-Whitney test | Uninjured ventral horn:<br>Ipsi vs Contra | 4, 4 | 7.0 ± 1.683 | 8.250 ± 3.092 | U = 6 | 0.6857 |
| Fig. 7F |  | Unpaired t-test | Ipsi caudal dorsal horn:<br>DMSO vs PAP4 | 10, 8 | 21.90 ± 4.233 | 42 ± 9.081 | t (16) = 2.147 | **0.0474** |
| Fig. 7G |  |  | Contra caudal dorsal horn:<br>DMSO vs PAP4 | 10, 8 | 11.33 ± 2.048 | 13.75 ± 3.385 | t (16) = 0.6271 | 0.5400 |
| Fig. 7K |  |  | Ipsi caudal intermediate gray:<br>DMSO vs PAP4 | 10, 8 | 6.200 ± 1.548 | 15.75 ± 3.509 | t (16) = 2.677 | **0.0165** |
| Fig. 7L |  |  | Contra caudal intermediate gray:<br>DMSO vs PAP4 | 10, 8 | 4.333 ± 0.9718 | 4.667 ± 0.3333 | t (16) = 0.2700 | 0.7914 |
| Fig. 7P |  |  | Ipsi caudal ventral horn:<br>DMSO vs PAP4 | 9, 8 | 1.667 ± 0.7638 | 7.375 ± 1.335 | t (15) = 3.820 | **0.0017** |
| Fig. 7Q |  |  | Contra caudal ventral horn:<br>DMSO vs PAP4 | 9, 8 | 0.7778 ± 0.3239 | 1.625 ± 0.5324 | t (15) = 1.395 | 0.1833 |
| Fig. 8B | Rostral PRV-positive cell counts | Unpaired t-test | Uninjured dorsal horn:<br>Ipsi vs Contra | 5, 5 | 61.20 ± 8.913 | 22.20 ± 2.035 | t (8) = 4.266 | **0.0027** |
|  |  |  | Uninjured intermediate gray:<br>Ipsi vs Contra | 5, 5 | 22.40 ± 4.632 | 17.20 ± 3.153 | t (8) = 0.9280 | 0.3806 |
|  |  |  | Uninjured ventral horn:<br>Ipsi vs Contra | 5, 5 | 7.0 ± 2.345 | 8.60 ± 2.088 | t (8) = 0.5095 | 0.6241 |
| Fig. 8F |  |  | Ipsi rostral dorsal horn:<br>DMSO vs PAP4 | 10, 8 | 1.9 ± 0.449 | 6.0 ± 1.822 | t (16) = 2.415 | **0.0280** |
| Fig. 8H |  |  | Ipsi rostral intermediate gray:<br>DMSO vs PAP4 | 10, 8 | 4.0 ± 1.366 | 4.625 ± 0.8438 | t (16) = 0.3656 | 0.7195 |
| Fig. 8I |  |  | Contra rostral intermediate gray:<br>DMSO vs PAP4 | 10, 8 | 5.5 ± 1.478 | 6.5 ± 0.7319 | t (16) = 0.5603 | 0.5830 |
| Fig. 8J |  |  | Ipsi rostral ventral horn:<br>DMSO vs PAP4 | 10, 8 | 2.80 ± 1.062 | 1.375 ± 0.3750 | t (16) = 1.148 | 0.2676 |
| Fig. 8K |  |  | Contra rostral ventral horn:<br>DMSO vs PAP4 | 10, 8 | 2.4 ± 0.8327 | 2.125 ± 0.6391 | t (16) = 0.2511 | 0.8049 |
| Fig. 8G |  | Mann-Whitney test | Contra rostral dorsal horn:<br>DMSO vs PAP4 | 10, 8 | 5.5 ± 1.928 | 4.250 ± 2.374 | U = 35 | 0.6756 |
| Fig. 9F | Syn+ puncta per PRV+ soma | Unpaired t-test | DMSO vs PAP4 | 5, 4 | 2.860 ± 0.7076 | 4.873 ± 0.2819 | t (7) = 2.397 | **0.0477** |
| Fig. 9G | 5HT+/syn+ puncta per PRV+ soma | Unpaired t-test | DMSO vs PAP4 | 5, 4 | 0.1906 ± 0.02820 | 0.2163 ± 0.05742 | t (7) = 0.4295 | 0.6805 |

**Supplemental table 4.**
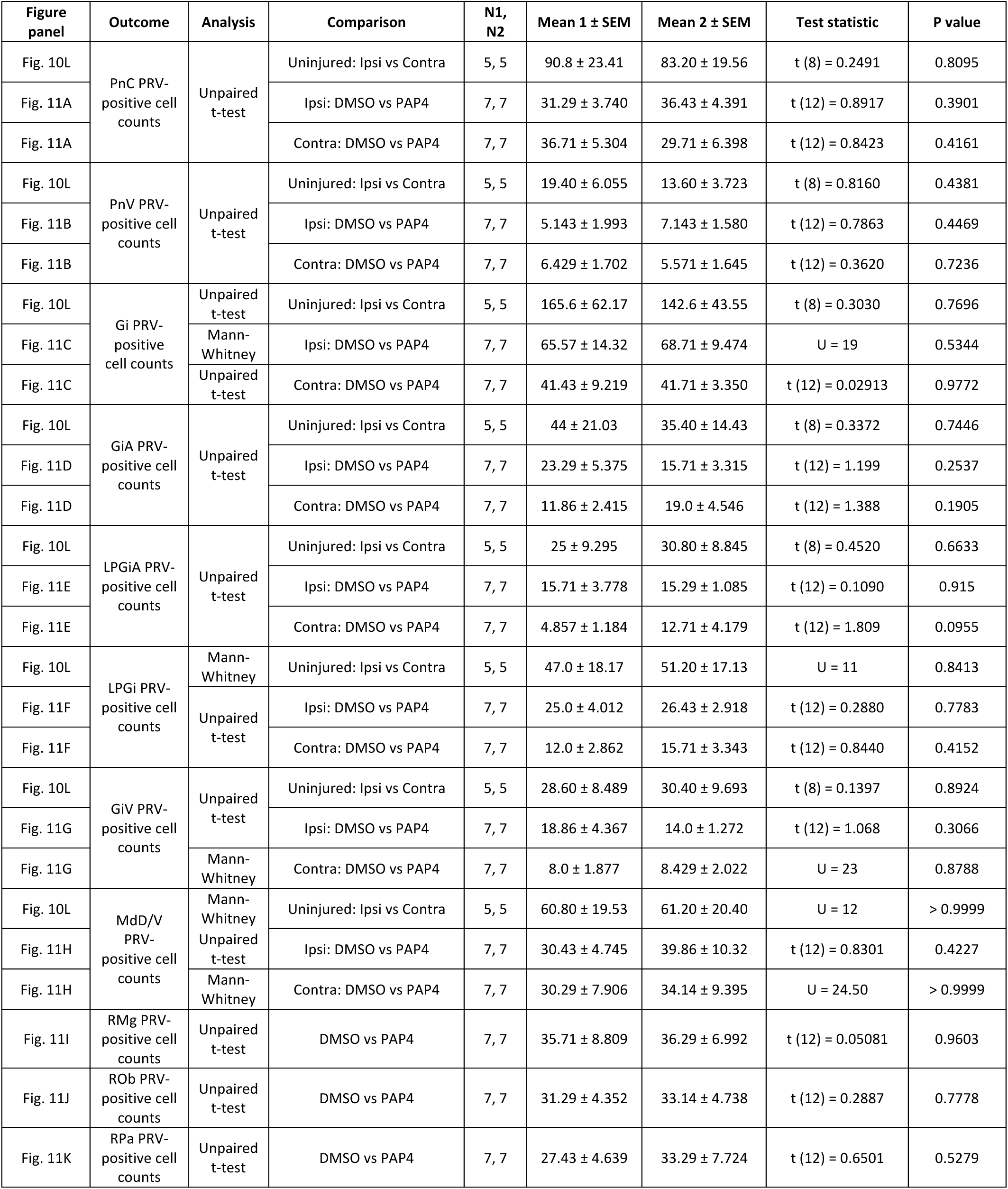

| **Figure panel** | **Outcome** | **Analysis** | **Comparison** | **N1, N2** | **Mean 1 ± SEM** | **Mean 2 ± SEM** | **Test statistic** | **P value** |
|---|---|---|---|---|---|---|---|---|
| Fig. 10L | PnC PRV-positive cell counts | Unpaired t-test | Uninjured: Ipsi vs Contra | 5, 5 | 90.8 ± 23.41 | 83.20 ± 19.56 | t (8) = 0.2491 | 0.8095 |
| Fig. 11A |  |  | Ipsi: DMSO vs PAP4 | 7, 7 | 31.29 ± 3.740 | 36.43 ± 4.391 | t (12) = 0.8917 | 0.3901 |
| Fig. 11A |  |  | Contra: DMSO vs PAP4 | 7, 7 | 36.71 ± 5.304 | 29.71 ± 6.398 | t (12) = 0.8423 | 0.4161 |
| Fig. 10L | PnV PRV-positive cell counts | Unpaired t-test | Uninjured: Ipsi vs Contra | 5, 5 | 19.40 ± 6.055 | 13.60 ± 3.723 | t (8) = 0.8160 | 0.4381 |
| Fig. 11B |  |  | Ipsi: DMSO vs PAP4 | 7, 7 | 5.143 ± 1.993 | 7.143 ± 1.580 | t (12) = 0.7863 | 0.4469 |
| Fig. 11B |  |  | Contra: DMSO vs PAP4 | 7, 7 | 6.429 ± 1.702 | 5.571 ± 1.645 | t (12) = 0.3620 | 0.7236 |
| Fig. 10L | Gi PRV-positive cell counts | Unpaired t-test | Uninjured: Ipsi vs Contra | 5, 5 | 165.6 ± 62.17 | 142.6 ± 43.55 | t (8) = 0.3030 | 0.7696 |
| Fig. 11C |  | Mann-Whitney | Ipsi: DMSO vs PAP4 | 7, 7 | 65.57 ± 14.32 | 68.71 ± 9.474 | U = 19 | 0.5344 |
| Fig. 11C |  | Unpaired t-test | Contra: DMSO vs PAP4 | 7, 7 | 41.43 ± 9.219 | 41.71 ± 3.350 | t (12) = 0.02913 | 0.9772 |
| Fig. 10L | GiA PRV-positive cell counts | Unpaired t-test | Uninjured: Ipsi vs Contra | 5, 5 | 44 ± 21.03 | 35.40 ± 14.43 | t (8) = 0.3372 | 0.7446 |
| Fig. 11D |  |  | Ipsi: DMSO vs PAP4 | 7, 7 | 23.29 ± 5.375 | 15.71 ± 3.315 | t (12) = 1.199 | 0.2537 |
| Fig. 11D |  |  | Contra: DMSO vs PAP4 | 7, 7 | 11.86 ± 2.415 | 19.0 ± 4.546 | t (12) = 1.388 | 0.1905 |
| Fig. 10L | LPGiA PRV-positive cell counts | Unpaired t-test | Uninjured: Ipsi vs Contra | 5, 5 | 25 ± 9.295 | 30.80 ± 8.845 | t (8) = 0.4520 | 0.6633 |
| Fig. 11E |  |  | Ipsi: DMSO vs PAP4 | 7, 7 | 15.71 ± 3.778 | 15.29 ± 1.085 | t (12) = 0.1090 | 0.915 |
| Fig. 11E |  |  | Contra: DMSO vs PAP4 | 7, 7 | 4.857 ± 1.184 | 12.71 ± 4.179 | t (12) = 1.809 | 0.0955 |
| Fig. 10L | LPGi PRV-positive cell counts | Mann-Whitney | Uninjured: Ipsi vs Contra | 5, 5 | 47.0 ± 18.17 | 51.20 ± 17.13 | U = 11 | 0.8413 |
| Fig. 11F |  | Unpaired t-test | Ipsi: DMSO vs PAP4 | 7, 7 | 25.0 ± 4.012 | 26.43 ± 2.918 | t (12) = 0.2880 | 0.7783 |
| Fig. 11F |  |  | Contra: DMSO vs PAP4 | 7, 7 | 12.0 ± 2.862 | 15.71 ± 3.343 | t (12) = 0.8440 | 0.4152 |
| Fig. 10L | GiV PRV-positive cell counts | Unpaired t-test | Uninjured: Ipsi vs Contra | 5, 5 | 28.60 ± 8.489 | 30.40 ± 9.693 | t (8) = 0.1397 | 0.8924 |
| Fig. 11G |  |  | Ipsi: DMSO vs PAP4 | 7, 7 | 18.86 ± 4.367 | 14.0 ± 1.272 | t (12) = 1.068 | 0.3066 |
| Fig. 11G |  | Mann-Whitney | Contra: DMSO vs PAP4 | 7, 7 | 8.0 ± 1.877 | 8.429 ± 2.022 | U = 23 | 0.8788 |
| Fig. 10L | MdD/V PRV-positive cell counts | Mann-Whitney | Uninjured: Ipsi vs Contra | 5, 5 | 60.80 ± 19.53 | 61.20 ± 20.40 | U = 12 | > 0.9999 |
| Fig. 11H |  | Unpaired t-test | Ipsi: DMSO vs PAP4 | 7, 7 | 30.43 ± 4.745 | 39.86 ± 10.32 | t (12) = 0.8301 | 0.4227 |
| Fig. 11H |  | Mann-Whitney | Contra: DMSO vs PAP4 | 7, 7 | 30.29 ± 7.906 | 34.14 ± 9.395 | U = 24.50 | > 0.9999 |
| Fig. 11I | RMg PRV-positive cell counts | Unpaired t-test | DMSO vs PAP4 | 7, 7 | 35.71 ± 8.809 | 36.29 ± 6.992 | t (12) = 0.05081 | 0.9603 |
| Fig. 11J | ROb PRV-positive cell counts | Unpaired t-test | DMSO vs PAP4 | 7, 7 | 31.29 ± 4.352 | 33.14 ± 4.738 | t (12) = 0.2887 | 0.7778 |
| Fig. 11K | RPa PRV-positive cell counts | Unpaired t-test | DMSO vs PAP4 | 7, 7 | 27.43 ± 4.639 | 33.29 ± 7.724 | t (12) = 0.6501 | 0.5279 |

